# Decoding *E. coli*’s Gut Survival Strategies: A CRISPRi Approach Across Diets, Inflammatory Environment and Strains

**DOI:** 10.64898/2026.01.05.697630

**Authors:** Amandine Maire, Emma Tkacz, Monica Ortelli, Christoph Dehio, Benoit Chassaing, Harry Sokol, Nathalie Rolhion, David Bikard

**Affiliations:** Institut Pasteur, Université Paris-Cité, UMR CNRS 3525, Synthetic Biology Laboratory, Paris, France; Sorbonne Université, Collège Doctoral, F-75005 Paris, France; Sorbonne Université, INSERM UMRS-938, Centre de Recherche Saint-Antoine, CRSA, AP-HP, F-75012 Paris, France; Gut, Liver & Microbiome Research (GLIMMER) FHU, F-75012 Paris, France; Biozentrum, University of Basel, Spitalstrasse 41, CH-4056 Basel, Switzerland; Institut Pasteur, Université Paris Cité, INSERM U1306, CNRS UMR6047, Microbiome-Host Interactions, Paris, France; Department of Gastroenterology, Saint-Antoine Hospital, Assistance Publique-Hôpitaux de Paris (AP-HP), F-75012 Paris, France; INRAE, UMR1319 Micalis, AgroParisTech, F-78350 Jouy-en-Josas, France

## Abstract

*Escherichia coli*, a ubiquitous member of the mammalian gut microbiota, exhibits remarkable genetic diversity underpinning its commensal or pathogenic lifestyles. Deciphering the precise genetic determinants enabling *E. coli*’s adaptation within the complex and dynamic intestinal environment is critical for understanding host-microbe symbiosis and enteric disease pathogenesis. Here, we establish an *in vivo* CRISPR interference (CRISPRi) platform that leverages bacterial gene fitness profiles as a high-resolution functional reporter to define the molecular niche and selective forces encountered by *E. coli* within mice harboring a defined minimal microbial community (OligoMM12). Our investigation revealed that dietary regimens profoundly reshape *E. coli*’s metabolic landscape and that the profile of essential genes help identify cross-feeding interactions. Comparative screens across a laboratory strain (MG1655), a Uropathogenic, and Adherent-Invasive *E. coli* (AIEC), identify distinct genetic requirements for intestinal colonization, highlighting divergent motility, stress response, and respiration strategies. In a host inflammatory environment, we find that the AIEC strain LF82 alters its colonization pattern, shifting towards the small intestine, and adapts to the inflammatory environment by remodeling its metabolism and stress responses. Notably, we uncover a critical role for mobile genetic elements, with the observation that inflammation triggers the induction of the Gally prophage which is beneficial for fitness in the healthy gut but becomes detrimental during inflammation. These findings provide a high-resolution genetic atlas of *E. coli*’s functional adaptation and demonstrate the utility of functional genomics to probe the gut environment itself.

## Introduction

*Escherichia coli*, a prevalent and genetically diverse member of the mammalian gut microbiota, can act as a harmless commensal or a versatile pathogen^1^. The various pathotypes of *E. coli* include intestinal pathogens such as Adherent-Invasive E. coli (AIEC), associated with Crohn’s disease, and extraintestinal pathogens such as Uropathogenic E. coli (UPEC) that may also have a commensal phenotype in the gut^2^. The ability of specific E. coli strains to colonize the gut or become pathogenic, depends not only on their genetic makeup but also on the complex ecosystem of the gut which is shaped by nutrient availability, host immune state, and inter-microbial interactions. Unravelling the genetic determinants that enable *E. coli* to adapt within these fluctuating intestinal niches is paramount for understanding symbiosis and enteric disease.

Functional genomics techniques, such as Transposon Insertion Sequencing (Tn-seq), have identified crucial bacterial fitness genes in animal hosts^3,4,5^, yet conducting such genome-wide studies *in vivo* presents significant technical hurdles. Tn-seq is limited by the need for massive mutant libraries, which are subject to *in vivo* population bottlenecks, as well as its inability to probe essential genes. While newer approaches like inducible Tn-seq^6^ or re-pooled arrayed libraries^7,8^ address some of these issues, their labor-intensive nature has restricted most studies to single strain and condition, limiting our understanding of adaptation to different environments modulated by diet and inflammation.

CRISPR interference (CRISPRi), which employs a nuclease-deficient Cas9 (dCas9) to silence target genes, has emerged as a transformative tool for high-throughput genomics. Its programmable and inducible nature allows the use of small guide RNA libraries well suited for *in vivo* experiments, while also enabling the study of genes that are essential under laboratory conditions. While CRISPRi screens have been successfully employed *in vitro*^9^, their adaptation for robust, genome-scale assessment of gene fitness directly within the animal gut remains to be achieved.

Here, using CRISPRi, we investigated *E. coli*’s adaptive strategies in the mammalian gut by exploring: (i) the impact of dietary regimens on the genetic requirements of commensal *E. coli* MG1655, (ii) strain-specific genetic requirements for gut colonization by comparing the commensal MG1655 to the adherent invasive *E. coli* (AIEC) LF82 and Uropathogenic *E. coli* (UPEC) CFT073 strains, and (iii) the adaptive responses of the AIEC strain LF82 to the host inflammatory environment in a DSS-induced colitis model, including the role of mobile genetic elements.

Collectively, these *in vivo* screens unveil previously unappreciated genetic pathways that govern *E. coli*’s survival, adaptation, and niche specialization. We identify specific metabolic pathways, stress responses, virulence factors, and mobile elements employed by *E. coli* to thrive under the diverse pressures of the mammalian gut, advancing our understanding of host-microbe interactions at a genetic level.

## Results

### Host diet shape E. coli gene essentiality profiles

We set up the CRISPRi screen in gnotobiotic C57Bl/6J mice carrying the OligoMM12 bacterial community^10^, providing a simplified and controlled mouse microbiota in which *E. coli* colonizes efficiently without the need for prior antibiotic treatment. Screens were performed using *E. coli* MG1655 carrying the EcoWG1 guide RNA library targeting all genes with an average of 5 guides per gene. Initial attempts to induce dCas9 expression in bacteria within the mouse by providing the inducer in the drinking water only yielded poor or no signal. Successful *in vivo* CRISPRi screens were ultimately achieved by pre-inducing dCas9 expression *in vitro* for 2 hours prior to gavaging of the *E. coli* strain carrying the guide RNA library, and then maintaining the inducer in the mice’s drinking water (Fig. S1 and Supplementary Text).

To study *E. coli* genetic adaptation to different gut environments, we fed OligoMM12 mice with different purified diets, with a known and controlled composition: a high-fat diet, a high-fiber diet, and a standard diet (Fig. 1A). A 16S analysis unsurprisingly revealed important shifts in microbiome composition between the 3 diets (Fig. 1B), which likely translate into differences in the niche available to *E. coli* and therefore differences in genetic requirements for efficient colonization. We next performed CRISPRi screens using *E. coli* MG1655 and sequenced the guide RNA library in the feces as well as in different sections of the intestines.

**Figure 1.**
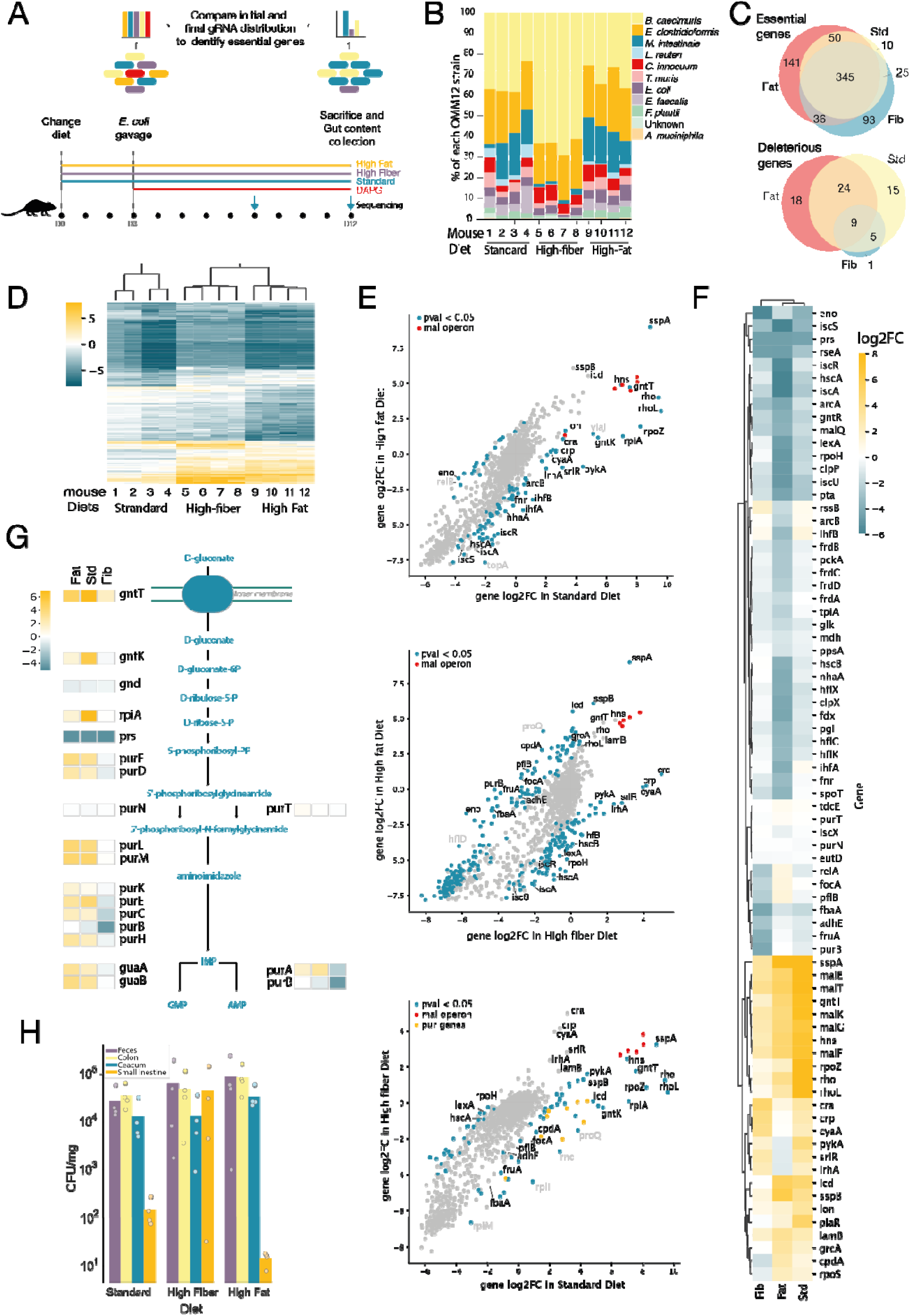
MG1655 gene essentiality in the gut under different dietary conditions. **(A)** Schematic of the experimental workflow for the *in vivo* CRISPRi screen comparing different dietary conditions. The *E.*LJ*coli* CRISPRi library was administered by gavage 3 days after the diet change. Sequencing was performed on feces 5 days post-gavage, and on both feces and different gut sections at the final time point. **(B)** 16S rRNA gene analysis of the OligoMM12 community composition in the feces of mice fed with different diets. **(C)** Venn diagrams showing the number of significantly essential (top diagram) or deleterious (bottom diagram) genes in each condition, identified by MAGeCK analysis. **(D)** the Heatmap showing hierarchical clustering based on the loglJ fold change (loglJFC) of genes with a significant difference in at least one pairwise comparison between diet conditions. **(E)** Pairwise comparisons of dietary conditions: scatter plots of gene loglJFCs between two diets. Blue dots indicate genes with a significant difference between conditions (FDR-corrected p-value < 0.05) and a median distance of all associated guides to the fitted line > 1. Red dots represent genes from the maltose operon. Specific genes discussed in the main text are labeled in black, other genes with a strong signal in grey. **(F)** Heatmap showing loglJFC values across the three dietary conditions for selected genes highlighted in the results section. **(G)** LoglJFC of genes involved in pathways ranging from gluconate transport to purine biosynthesis, across the three dietary conditions. **(H)** *E. coli* colonization levels in different gut regions, expressed as CFU per mg of content.

A total of 700 genes were identified as essential, and 72 as deleterious, in at least one condition. Among them, 244 essential and 35 deleterious genes showed a significant effect in only one specific diet (Fig. 1C, FigS2). A hierarchical clustering analysis performed on gene effects at D8 revealed that replicates from the same dietary condition cluster together, indicating clear differences in the set of genes used by *E. coli* to adapt to the different gut environments. Conversely, the results obtained from a given feces sample can be used to easily infer the diet received by the animal, suggesting that CRISPRi screens can be used to investigate the state of the gut environment, using *E. coli* as a “sentinel cell”, in a highly reproducible manner (Fig. 1D).

### Deleterious genes reveal the misadaptation of MG1655 to the gut environment

First, we observed that the repression of some genes by dCas9 enhances the fitness of *E. coli* across all dietary conditions, suggesting these genes are deleterious in the animal gut, and that MG1655 is not well adapted to this environment. In contrast, strongly deleterious genes cannot be identified from *in vitro* screens performed in LB^11^ (Fig. S3C), likely due to the laboratory adaptation of *E. coli* MG1655 and the relative lack of selective pressure (Fig. 1EF). Key genes found to be detrimental in the gut included maltose transporters (*malEFG* and *malK*-*lamB* operons), and gluconate transporter (*gntT*). Additionally, silencing several global regulators such as *hns*, *sspA*, *sspB* and the transcription terminator *rho*, also enhanced fitness (see Supplementary Text for more details). These results suggest that MG1655 fails to properly adjust its gene expression for *in vivo* growth, and as a result, silencing certain deleterious genes or master regulators with dCas9 can facilitate its survival.

### Diet shapes E. coli metabolic niche

We next investigated to which extent different diets could reshape *E. coli*’s metabolic gene dependencies. In standard and high-fiber diets, genes involved in cAMP-CRP complex formation (*cyaA*, *crp*) were found to be detrimental. cAMP activates the expression of catabolic genes in the absence of rapidly metabolizable carbon sources. This likely reflects the presence of glucose derived from starch degradation by host or bacterial amylases in the standard and high-fiber diets. In addition, the fructose transporter *fruA* becomes important to *E. coli* in the high-fiber diet only. This diet contains a large amount of inulin which might be released as fructose by inulinase-producing members of the OligoMM12 community, notably *Bacteroides caecimuris*, whose relative abundance in the gut showed a marked increase in this diet (Fig. 1B). Other signals in the high-fiber diet are consistent with these observations, including the deleterious effect of *cra*, which was previously shown to repress the fructose operon and enhance *cyaA* activity^12^, and the beneficial effect of *cpdA*, which degrades cAMP. In contrast, in the high-fat diet, the “low glucose/fructose” environment likely increases the importance of gluconeogenesis which is activated by cAMP-CRP. This hypothesis is supported by the observed importance of genes like *pckA*, *pgi*, *tpiA*, *ppsA*, and *mdh*, in this diet.

Another strong signal for genes involved in carbohydrate metabolism is the positive effect of silencing the repressor of the sorbitol operon *srlR,* thereby activating the sorbitol utilisation operon in diets containing starch (standard and high-fiber diets), suggesting that sorbitol, which is absent from the diet, is produced by other members of the OligoMM12 community (Supplementary Text).

The effect of diet can further be detected in core metabolic pathways including glycolysis (*pgi*, *eno*, *fbaA*, *pykA*) and mixed acid fermentation (*eutD*, *adhE*, *pta*). Of note, in the high-fat diet, *E. coli* seems to favor fermentation pathways that produce succinate (*frdABCD)* over those that produce formate *(grcA, tdcE* and *focA-pflB* operon) and oxoglutarate (*icd*). A possible explanation is that the production of oxoglutarate and the oxidation of formate waste carbon in the form of CO_2_^13^, while fermentation towards succinate does not, which might be beneficial in a low carbohydrate environment.

A particularly strong signal is the deleterious effect of genes involved in *de novo* synthesis of AMP and GMP from gluconate in standard and high-fat diets. This includes 14 steps from the import of gluconate, going through gluconate degradation, PRPP biosynthesis, 5-aminoimidazole ribonucleotide biosynthesis, inosine-5’-phosphate biosynthesis and finally guanosine and adenosine biosynthesis. The only exceptions being *gnd* and the expected essential genes (*purB*, *prs*) and redundant genes (*purN*, *purT*) (Fig. 1G). The standard and high-fat diet environments might be richer in purines than the high-fiber diet, rendering *de novo* synthesis unnecessary and costly. Silencing this pathway might also help maintain a balanced nucleotide pool.

The high cost of expressing the gluconate transporter *gntT* further suggests that the gut environment in standard and high-fat diets is richer in gluconate. Previous work suggests that gluconate is readily available in the host mouse mucus^14^, but gluconate could also be formed by other members of the microbiome as a product of glucose-fructose oxidoreduction^15^, alongside sorbitol, or through glucose oxidation by glucose dehydrogenases^16,17^.

#### High-fiber diets create a more favourable environment for E. coli in the small intestine

The abundance of *E. coli* in each region of the gut (small intestine (SI), cecum, colon, and feces) was evaluated at the end of the experiment. This revealed different colonization patterns across diets, specifically in the SI (Fig. 1H). While this region is typically less colonized by *E. coli*, a high-fiber diet seems to promote a bloom in the SI, supporting the idea that *E. coli* colonization patterns can be influenced by diet^18^. Furthermore, comparison of *E. coli* gene essentiality profiles in the SI across the three dietary conditions showed a distinct profile for the high fiber diet. Notably, genes involved in fructose metabolism (*fruAKB*) and the multidrug efflux pump (*acrAB*) were essential in the SI of high-fiber-fed mice (Fig S4A-C). *AcrAB* is known to contribute to resistance to bile acids which are abundant in the SI, and was previously identified as a colonization factor of Enterohaemorrhagic *E. coli* in the infant rabbit gut^4^. In contrast, genes involved in iron-sulfur (Fe-S) cluster biogenesis were deleterious under the high-fiber diet but essential in the other diets (*fdx*, *hscB*) (Fig S4AC).

When comparing gene essentiality profiles between fecal and SI *E. coli* populations within the high-fiber condition, we also observed region-specific differences (Fig S4D). For example, genes involved in translation (*ssrA*, *glnV*), fatty acid biosynthesis (*fabH*), and the stringent response (*dksA*) were essential in feces but deleterious in the SI. Altogether, these findings demonstrate that dietary composition influences both the spatial colonization patterns of *E. coli* and its genetic requirement to adapt to different gut regions.

### Strain-specific gut adaptation

We next decided to investigate how different *E. coli* strains adapt to the gut by comparing the laboratory strain MG1655 with two clinical strains: the AIEC LF82 and the uropathogenic CFT073. Using whole-genome CRISPRi libraries customized for each strain, we performed parallel screens in OligoMM12 mice that were fed a standard chow diet (Fig. 2A). Significant essential and deleterious genes at the end point (D9) have been identified in each strain (Fig. S5A, Fig. S6). The clustered heatmap of those gene essentiality profiles revealed that the two clinical strains are more closely related to each other than to the laboratory strain (Fig. 2B). This suggests that LF82 and CFT073, both from the *E. coli* phylogroup B2 , may use similar survival strategies to adapt to the gut environment, distinct from MG1655. These findings are consistent with the observations that both pathogenic strains colonized the gut more effectively than MG1655 (Fig. S7).

**Figure 2.**
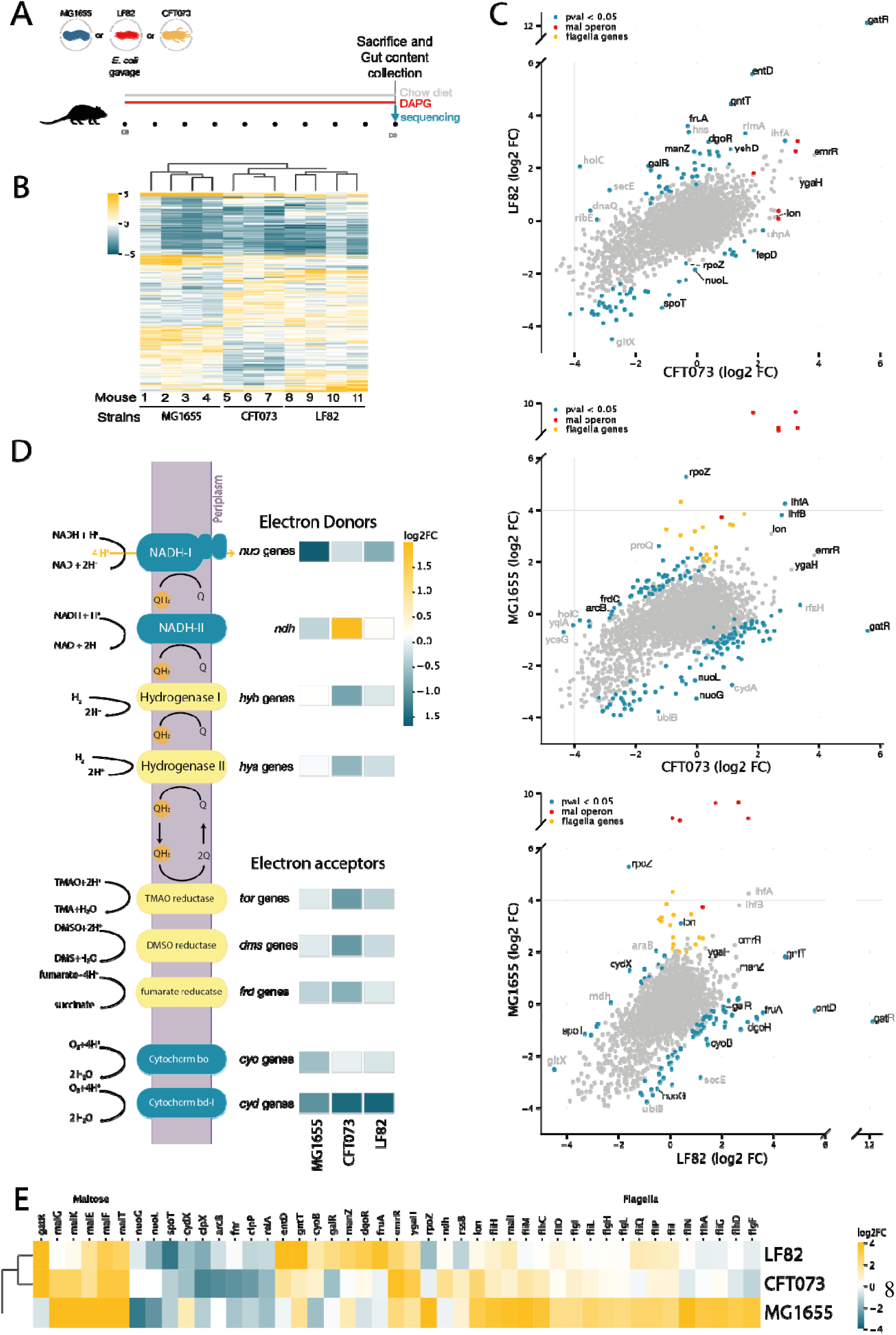
Comparison of gene essentiality in MG1655, LF82, and CFT073 *E. coli* strains in the gut. **(A)** Schematic of the experimental workflow for the *in vivo* CRISPRi screen comparing different *E. coli* strains. **(B)** Heatmap showing hierarchical clustering based on the loglJ fold change (loglJFC) of genes with a significant difference in at least one pairwise comparison between strains. **(C)** Pairwise comparisons of strains: scatter plots of gene loglJFCs between two strains. Blue dots indicate genes with a significant difference between strains (FDR-corrected p-value < 0.05) and a median distance of all associated guides to the fitted line > 1. Red and purple dots represent genes from the maltose operon (*mal*) and flagellar operons (*fli*, *flg*, *flh*), respectively, that are significantly different between the strains. Labeled genes are those highlighted in the results section and show a strong differential signal. Specific genes discussed in the main text are labeled in black, other genes with a strong signal in grey. **(D)** LoglJFC of operon involved in electron transport chain activity across the three strains, highlighting differences in the use of electron donors and acceptors involved in aerobic (blue) and anaerobic (yellow) respiration. For each complex protein, the median log2FC of all guides across all the genes of the operon is calculated. **(E)** Heatmap showing loglJFC values across the three strains for selected genes discussed in the results section.

The distinct clustering of MG1655 partly reflects the presence of genes that are deleterious for this strain, and which are less detrimental or even neutral for the clinical strains. This includes genes involved in maltose transport and their activator (*malT*), as described above, as well as genes involved in flagellar protein expression and assembly (*flg*, *fli*, *flh* operons) (Fig. 2C,E) (see Supplementary Text). While not as strong as for MG1655, the clinical strains also showed signs of misadaptation to the artificial setup of the OMM12 mouse gut. This included the beneficial effect of silencing the efflux pump repressor *emrR*, or the repressor of the galactitol utilization pathway *gatR*. The strong signal observed on *gatR* suggests that LF82 and CFT073 fail to de-repress the *gat* operon, even though they would benefit from metabolizing galactitol. In MG1655, *gatR* is broken due to an insertion of IS*3E*, which likely explains why we do not see any signal for this strain. (see Supplementary Text)

Focusing on genes essential in some strains and not others provides insights into the distinct environmental niches and stresses they face. Silencing SpoT, which both synthesizes and hydrolyzes (p)ppGpp, a key regulator of the stringent response, affects LF82’s fitness but is neutral in MG1655 and CFT073. Conversely, *lon* and *rpoZ*, which mediate stress adaptation by degrading misfolded proteins (Lon) and modulating ppGpp-dependent transcription (RpoZ), are deleterious to MG1655 in chow and standard purified diets, yet neutral for the other strains. Stress response requirements thus appear to be both strain- and diet-dependent. Supporting this, in the high-fat diet, the stringent response seems detrimental to MG1655: silencing the (p)ppGpp synthetase *relA* improves fitness, while silencing its hydrolase *spoT* reduces it. Similarly, silencing *rpoS* boosts fitness, while silencing *rssB/clpXP* which degrades *rpoS* is detrimental.

Genes involved in respiration display significant differences in essentiality across the three *E. coli* strains and suggest that CFT073 inhabits a more anaerobic niche than MG1655 and LF82 (see Fig. 2D and Supplementary Text). Consistently the *frd*, *tor*, and *dms* genes, which encode alternative reductases that use fumarate, TMAO, or DMSO respectively as terminal electron acceptors under anaerobic conditions, appear more important in CFT073. The importance of *arcB* and *fnr* which activate and regulate genes involved in anaerobic respiration, further supports this hypothesis.

Pathobionts carry virulence factors that can facilitate their colonization and invasion. We looked at the impact of genes that could contribute to the higher colonization level of LF82 and CFT073 compared to MG1655. Iron is a known limiting resource for bacterial growth in the gut environment and enhanced iron uptake is a known virulence factor. Silencing *entD*, which is involved in the synthesis of the enterobactin siderophore, gives a strong fitness advantage to both CFT073 and LF82 *in vivo* but not *in vitro*. The signal observed suggests that for these strains, enterobactin biosynthesis is highly active and thus very costly. Transcriptomics data from a previous study has indeed shown that siderophore synthesis and transport are upregulated *in vivo* in an AIEC strain^19^. Bacteria that silence *entD* however likely benefit from enterobactin produced by neighboring cells, or from other iron acquisition systems, which could explain their fitness advantage. Further corroborating the low iron availability in the gut environment, we observed that microcin MccH47 carried by CFT073 is essential in the gut but neutral *in vitro* (Fig. S3B). The expression of this microcin is known to depend on iron availability and increases under iron-limiting conditions^20^.

Genes involved in capsule synthesis are also known to play an important role in colonization and immune evasion. In CFT073, *kpsC*, a gene part of the *kps* operon involved in assembly of group 2 capsular polysaccharides (K antigen), is slightly deleterious *in vitro* but highly essential *in vivo* (Fig. S3B). This echoes the previously reported importance of the K1 capsule for E. coli colonization of the neonatal rat intestine^5^. KpsC is absent in MG1655 but present in LF82 where it shows a similar yet weaker signal. These findings suggest that the capsule of CFT073 and LF82 contribute to their efficient colonization of the mouse gut.

### LF82 adaptation to the inflamed gut environment

We further leveraged our method to investigate how inflammatory conditions affect the genetic requirements for colonization in both MG1655 and the Crohn’s disease associated strain LF82. To this end, we conducted CRISPRi screens in a DSS-induced colitis mouse model. Mice were treated with DSS in the drinking water for 7 days to induce colitis. On day 7, corresponding to the peak of inflammation, DSS was replaced with water supplemented with DAPG, mice were orally gavaged with either MG1655 or LF82 and feces sampled over a 8 days period (Fig. 3A).

**Figure 3.**
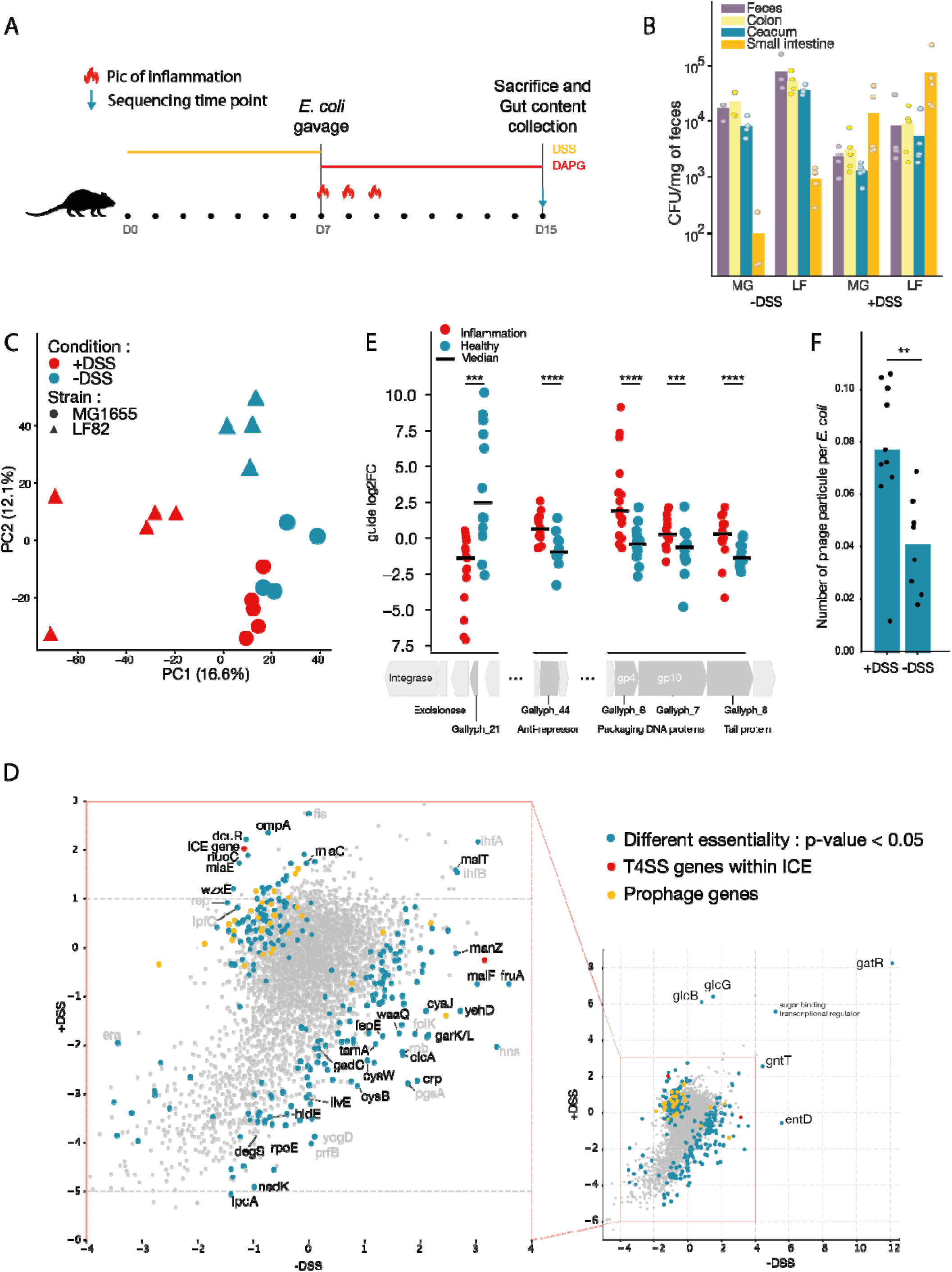
LF82 genetic adaptation to the inflamed gut. **(A)** Schematic of the experimental workflow for the *in vivo* CRISPRi screen in the DSS-induced colitis model. **(B)** LF82 colonization levels in different gut regions in HlJO- and DSS-treated mice, expressed as CFU per mg of content. **(C)** Principal component analysis (PCA) based on gene loglJ fold change (loglJFC) values from MG1655 (circles) and LF82 (triangles) in HlJO-treated (blue) or DSS-treated (red) mice. **(D)** Scatter plot showing gene loglJFC values of LF82 genes in HlJO- versus DSS-treated mice. Blue dots indicate genes with a significant difference between conditions (FDR-corrected p-value < 0.005) and a median distance of all associated guides to the fitted line > 1. Among them, yellow and red dots correspond to genes from prophages and integrative conjugative elements (ICEs), respectively. Specific genes discussed in the main text are labeled in black, other genes with a strong signal in grey. **(E)** LoglJFC values of guides targeting the Gally genome, focusing on genes (dark grey) that show strong differential signals between *H*LJ*O-* (blue) and DSS-treated (red) mice. Each dot represents an individual guide in one mouse. Statistical significance (FDR-adjusted): ****□p□≤□0.0001, ***□p□≤□0.001. **(F)** Quantification of Gally phage particle abundance in feces of *H*LJ*O-* and DSS-treated mice. qPCR targeting the Gally genome was performed on fecal supernatants and normalized to both qPCR results targeting the LF82 genome from the same samples to control for genomic DNA contamination. The number of Gally particles was normalized to the total number of *E. coli* in the pellet from the same fecal sample. Statistical significance: **□p_value□≤□0.01.

The 16S rRNA sequencing data showed that inflammation does not significantly affect the relative abundance of each species in the gut (fig. S8A). However, LF82 appeared to alter its colonization pattern, shifting toward the SI (Fig. 3B). Interestingly, this shift recapitulates AIEC blooms in the ileum described in Crohn’s disease patients ^21^. A principal component analysis of CRISPRi screening data from MG1655 and LF82 revealed a clear separation between healthy and inflamed environments, further highlighting how CRISPRi screens can effectively report on the state of the gut environment. The gene fitness profile of LF82 shifted more dramatically than MG1655’s in response to inflammation, indicating a greater adaptation (Fig. 3C). We therefore focused on analyzing the strategies LF82 uses to survive in the inflamed gut. Genes that are significantly essential or deleterious for LF82 under both +DSS and –DSS conditions were identified from end point sequencing (Fig. S5B, Fig. S9).

The inflamed gut has been found to exhibit increased oxidative stress^22^ , lower pH^23^, iron scarcity^24,25,26,27^, and higher oxygen^28^. Consistently, genes involved in acid tolerance (*clcA*), and high-affinity iron uptake (*entD*, *fep* operon) became more critical to LF82, while the activator of anaerobic fumarate respiration (*dcuR*) became deleterious in the inflamed gut (Fig. 3D). Several signals also point to an impaired redox balance and oxidative stress during inflammation. The NDH-I complex (*nuoC*), which consumes NADH, became deleterious during inflammation, while NADH-generating *lpdA* (TCA cycle) and NADP-generating *nadK* became strongly essential. Interestingly, one of the strongest signals during inflammation is the deleterious effect of *glcB* (and the upstream *glcG*), which produces malate, a key metabolite of the TCA cycle^29^. We hypothesise that LF82 expresses *glcB* to maladaptive levels, disrupting the TCA cycle, and that silencing it helps restore the NAD(P)H pool, which is crucial to maintain a reduced environment and defend against oxidative stress^30^. This is further supported by the increased importance of sulfate assimilation genes (cysJ, cysW, cysB), which are key for synthesizing antioxidants like hydrogen sulfide (H2S)^31^, cysteine and glutathione.

### Inflammation drives a shift in colonization and metabolic strategy

LF82 shows an important shift in sugar metabolism during inflammation with an increased importance of *crp*, while *fruA*, *gntT*, *manZ* and *malF* which were deleterious in healthy condition became neutral. In addition, genes involved in the use of galactarate (*garK*, *garL)* became important, indicating an increase in galactarate availability. Interestingly, reactive nitrogen and oxygen species were described to non-enzymatically oxidize dietary or host-derived sugars, including galactose to galactarate^32^. A similar reprogramming was observed for amino acids. The *de novo* synthesis of branched-chain amino acids via the *ilv* genes became important (*ilvE*). Concurrently, the glutamate transporter *gadC* became essential, suggesting that glutamate remains accessible.

### The envelope stress response becomes important during inflammation

The inflamed environment appears to generate increased envelope stress in LF82. The sigma factor gene *rpoE,* and genes involved in LPS biosynthesis like *waaQ* and *hldE*, are more important in DSS condition than in the healthy gut. Consistently, *degS*, coding for a membrane-anchored protease that senses misfolded OMPs and activates *rpoE,* also became more important. LF82 further benefited from silencing *ompA* during inflammation, a gene known to be repressed by *rpoE* (Supplementary Text). Furthermore, silencing the Enterobacterial Common Antigen (ECA, *wzxE*) favored LF82’s colonization, possibly by making it less visible to the host immune system.

### Mobile genetic element involved in LF82’s fitness in healthy and inflamed gut

Among the genes exhibiting significant differences between inflamed and healthy conditions, many are located within LF82’s prophages (Fig. 3D), and some within its integrative and conjugative elements (ICEs). This suggests that mobile genetic elements contribute to adaptation to the inflamed gut environment.

LF82 harbors an ICE with two genes, located in the type IV secretion system (T4SS), that display a strong differential signal between the healthy and inflamed conditions (Fig. S10). Genes belonging to the T4SS operon generally appeared to be more deleterious during inflammation, while a gene in a nearby operon showed the opposite signal, which could indicate that it acts as a regulator of the T4SS. The T4SS has previously been identified as a key colonization factor for AIEC and is known to be activated by host signals to promote biofilm formation^19^.

Prophage induction also appeared to affect colonization differently in the healthy versus inflamed gut. LF82 contains four prophages (Gally, Perceval, Tritos and Cartapus)^33^. As expected, repressing the prophage repressors, thereby inducing their lytic cycle, was detrimental to LF82 across conditions and phages. This indicates that those prophages can undergo lytic induction *in vivo*, which was previously undocumented for Cartapus (fig. S10). We further observed that under inflammatory conditions, blocking the lytic cycle of Gally by repressing the anti-repressor or other genes of the lytic operon confers a fitness advantage. Conversely, a gene (Gallyph_21 in GenBank OV696608.1) located near the excisionase and integrase exhibits the opposite signal, suggesting that it plays a role in the maintenance of lysogeny (Fig. 3E). These signals suggest an increased lytic activity of the phage in the inflamed gut to levels that are detrimental for the colonization of LF82. Measuring the abundance of Gally particles in the feces of inflamed versus healthy mice indeed showed a significant 2-fold increase during inflammation (Fig. 3F). Surprisingly, silencing Gallyph_21 in healthy condition was even beneficial to *E. coli*, suggesting that LF82 benefits from an increased level of prophage induction, likely not proceeding to a full lytic cycle, in the healthy gut. Together, these results highlight a complex relationship between Gally and its host that is shaped by the environment.

## Discussion

In this study, we report that *E. coli’s* genetic requirements for gut colonization vary significantly both in between strains, and as a function of the gut environment, such as those shaped by different dietary regimens or inflammation. Our results provide an atlas of genes involved in adaptation to the gut, with a total of 767 genes found to have a significant effect in at least one environment. Our data show that *E. coli*, and the laboratory adapted MG1655 in particular, fails to properly regulate key systems like maltose import and the flagellum. Consistently, previous studies of *E. coli* adaptation to the murine gut identified frequent mutations in these genes^34^. Overall, the unique pattern of gene essentiality provides a rich signature of the environment, establishing that *in vivo* CRISPRi screening can serve as a functional probe to assess the state of the gut.

While the OligoMM12 model does not recapitulate all the processes of a complex microbiome^35^, it offers a highly interpretable and well-controlled experimental setup to investigate adaptation strategies, especially when paired with purified diets. Our results reveal that *E. coli* employs distinct metabolic strategies that reflect nutrient availability, supporting the "restaurant hypothesis" where different microbial niches offer unique "menus" via cross-feeding^36^. This is highlighted in the high-fiber diet, where fructose utilization became important despite the absence of fructose in this diet. This shift coincided with increased colonization by *B. caecimuris*, which is expected to release fructose as a result of inulin digestion.

Among our strongest signals was that gluconate utilization, previously reported as beneficial to *E. coli* in the gut environment^14,37,38^, is highly deleterious under high-fat and standard purified diets. The observation of the same signal for all downstream genes from gluconate import to the synthesis of GMP and AMP, suggests that these diets lead to a costly overproduction of purines by MG1655.

Experiments with different *E. coli* strains revealed diverse adaptation strategies. Pathogenic strains colonized to a higher level and relied more on genes described as virulence factors. This includes iron uptake systems, capsule biosynthesis and type IV secretion system. Strains also seem to occupy distinct respiratory micro-niches, with the uropathogenic CFT073 in particular relying more on anaerobic respiration.

Focusing on the AIEC strain LF82 in an inflamed gut, we found that it adapts by shifting its metabolism, and by responding to oxidative and envelope stress. Mobile genetic elements like the Gally prophage are also involved in adaptation. We found that its lytic cycle is induced to levels that impact bacterial fitness in the inflamed gut, which might result from a more genotoxic environment. Interestingly, induction of Gally was recently shown to be suppressed in macrophages^33^, in which LF82 also undergoes genotoxic stress, suggesting that the presence or absence of other signals are needed to drive prophage induction. Intriguingly, silencing a gene upstream of the excisionase appears beneficial in healthy conditions. A possible explanation might be that the excision of this prophage doesn’t always lead to bacterial death and may regulate chromosomal genes essential for gut survival, a phenomenon previously described for several phages and species^39–42^. The Gally prophage integrates in front of the *torS* gene, the sensor of the torR/S system involved in the regulation of the *torCAD* operon which encodes a trimethylamine N-oxide (TMAO) reductase^43^. Previous work on another phage, HK022, showed that its integration at this exact locus shuts-off the TMAO reductase under aerobic conditions^44^. We therefore hypothesize that the excision of Gally in the healthy gut could similarly modulate the *tor* system, fine-tuning TMAO respiration in a way that promotes colonization.

In conclusion, our work reveals the remarkable plasticity of *E. coli*, showing that its genetic requirements for gut colonization are dynamically dictated by the host and microbial environment. This functional genomics approach provides a detailed atlas of bacterial adaptation and establishes a novel framework for using bacterial CRISPRi screens as sensitive probes to assess the state of the gut.

## Materials and Methods

### Bacterial strains and media

The CRISPRi screen was performed on three *Escherichia coli* strains: MG1655, LF82, and CFT073. All strains were rendered streptomycin-resistant via oligo recombineering using the pKOBEG plasmid, following the protocol described by Chaveroche et al.^45^. Specifically, the K43R mutation in the *rpsL* gene was introduced using the AM283 primer (Table S1).

*E. coli* K-12 strains MG1655Δ*fhuA* and MFDpir^46^ were used for cloning. Cultures were performed in Luria-Bertani (LB). When mentioned chloramphenicol (Cm) was used at 20 µg/mL, kanamycin (Kan) at 50 µg/mL, diaminopimelic acid (DAP) at 0.3 mM, and 2,4-Diacetylphloroglucinol (DAPG) at 25 µM, streptomycin at 100µg/mL, and CaCl2 at 5mM. Drigalski Agar medium was used to select *E. coli* from stool or organ samples.

### CRISRPi library design

***MG1655***: The CRISPRi library used to target the MG1655 genome was the EcoWG1 library developed by Calvo-Villamán et al^47^. This library comprises approximately 20,000 sgRNAs, with five guides per gene.

***LF82 and CFT073***: Genome-wide CRISPRi libraries for LF82 and CFT073 (LF82WG and CFT073WG) were constructed by combining the EcoCG library, which targets the *E. coli* core genome and contains 11,629 sgRNAs (three guides per gene), with custom add-on libraries designed to target each strain’s accessory genome. Add-on libraries were built using the *generate_library.py* script, which first identifies all possible sgRNAs in the target genome containing a protospacer adjacent motif (PAM) within coding regions. For each locus tag covered by fewer than three guides in the EcoCG library, one to three additional sgRNAs were selected from this list to ensure uniform coverage (three guides per gene, when possible).

The additional guides were selected according to the design rules outlined above, prioritized in the following order : (i) sgRNAs with unique targets in the genome, (ii) minimal off-targets with perfect identity to the 12-nt seed region (noff_12) (iii) minimal off-targets on the non-template strand with 11-nt seed identity (noff_11_gene) (iv) minimal off-targets in promoter regions with 9-nt seed identity (noff_9_prom) (v) predicted activity scores (per the Calvo-Villamán et al. model)C(iv) targets located in the first half of the gene. Each of the LF82 and CFT073 add-on libraries included ∼4,000 sgRNAs. Additionally, approximately 2000 sgRNAs from the EcoCG library did not match any target in the LF82 or CFT073 genome, they were included as non-targeting controls guides.

### CRISPRi library construction and transformation

All the CRISPRi Libraries were cloned on the pFR56 plasmid^9^ This plasmid carries dCas9 under the control of the DAPG inducible pPhlF promoter. Additionally, it includes two BsaI sites for cloning sgRNA guides via Golden Gate assembly, enabling their expression under a constitutive promoter.

### Conjugation of EcoWG1 library in streptomycin resistant MG1655

For the *E. coli* MG1655 strain, the EcoWG1 CRISPRi library was stored in the conjugative donor strain MFDpir::pFR58, which harbors the plasmid pFR58. This plasmid, which encodes PhlF protein repressing dCas9 promoter, provides an additional dCas9 repression in order to prevent leaky expression of dCas9 and limit the introduction of potential biases in the library. Conjugation into streptomycin-resistant MG1655 was performed as follows: A 1:50 pre-culture of the donor strain (MFDpir::pFR58 + EcoWG1) was prepared by inoculating an entire glycerol stock into 50 mL of LB supplemented with DAP, Kan, and Cm. In parallel, a 1:50 culture of streptomycin-resistant MG1655 was prepared in LB from an overnight pre-culture. When both donor and recipient cultures reached an ODCCC of 0.6, cells were washed to remove antibiotics. Equal volumes (2 mL) of donor and recipient cultures were mixed, centrifuged, and resuspended in 100 µL of LB. The entire volume was spotted onto an LB+DAP plate and incubated at 37°C for 2 hours to allow conjugation. Following incubation, the bacterial spot was scraped with an inoculating loop, resuspended in 1 mL of LB, and 300 µL was plated onto 3 square LB+Cm plates (900 µL total). An additional 100 µL was used for serial dilutions and plated on LB+Cm to estimate conjugation efficiency. Donor (MFDpir) and recipient (MG1655) strains were plated on LB+Cm as negative controls. After only 4 hours of incubation at 37°C (to avoid overgrowth of the colonies and limit potential biases in library composition), transconjugants were recovered by washing each plate twice with 2.5 mL of LB. The resulting 15 mL were pooled and aliquoted into 15 glycerol stocks for storage. This procedure yielded approximately 10C transconjugants, representing a ∼50-fold coverage of the library.

### Cloning and transformation of the LF82WG and CFT073WG guide RNA libraries

The guides from the custom-designed add-on_LF82 and add-on_CFT073 libraries were synthesized by Twist Bioscience, in the following format: 5′-AGCTGTCACAGGTCTCATAGT-[guide_sequence]-GTTTTGAGACCGCGACTACGTCTAC-3′. For each add-on library, six PCR reactions were performed on 0.625 ng of the single-stranded oligo pool using primers Sol70 and Sol71 (Table S1) to generate the second DNA strand and amplify the pool. PCR reactions were done with KAPA HiFi polymerase (Roche) and PCR conditions were as follow; initial denaturation at 95 °C for 3 min; six cycles of 98 °C for 20Csec, 57 °C for 15Csec, and 72 °C for 15Csec; followed by a final extension at 72 °C for 1Cmin. The six PCR products were pooled and purified using a PCR clean-up column (Macherey-Nagel), eluted in 20 µL of nuclease-free water, yielding 2.35 ng/µL of double-stranded DNA. The resulting library was cloned into the pFR56 plasmid via Golden Gate assembly using ∼8 nM of plasmid and ∼24 nM of insert (1:3 molar ratio). The assembly reaction was drop-dialyzed for 20 minutes and transformed into the CH-E02 strain, an *E. coli* MG1655Δ*fhuA* strain carrying a chromosomally integrated acrIIA4 anti-CRISPR gene. This anti-CRISPR prevents dCas9 activity in case of leaky expression, ensuring that no bias is introduced in the library during cloning. Electrocompetent CH-E02 cells were prepared from a culture stopped at OD₆₀₀ ≈ 0.7 and washed three times with 10% glycerol. The final pellet was resuspended in water to concentrate the cells approximately 2,000-fold. Two electroporations were performed using 80 µL of cells and 5 µL of the Golden Gate reaction mixture each. After a 1-hour recovery at 37°C, cells were plated on square LB+Cm plates (100 µL per plate) and incubated overnight at room temperature. To assess transformation efficiency, serial dilutions of the recovered culture were plated on LB+Cm and incubated overnight at 37°C. The transformation yielded sufficient coverage, exceeding 100-fold of the library. The next day, plates were washed three times with LB, and glycerol stocks were prepared for long-term storage at –80°C. To obtain each of the LF82WG and CFT073WG libraries, plasmid minipreps of the respective add-on and EcoCG libraries were performed from cultures inoculated 1:20 from glycerol stocks and grown for 3 hours at 37°C in LB+Cm. The two libraries were then combined at a 1:2.5 ratio (add-on : EcoCG), adjusted based on their respective sizes. A total of ∼100 ng of the mixed library was electroporated into 100 µL of electrocompetent, streptomycin-resistant LF82 or CFT073 cells (prepared as described above). Selection and storage followed the same procedure as described for the CH-E02 transformation. After overnight incubation at room temperature, plates were washed twice with LB, and glycerol stocks of the transformants were prepared and stored at –80°C.

### Mice

All experiments were conducted using gnotobiotic OligoMM12 mice^10^, bred at the Institut Pasteur (Paris, France). Animals were maintained in isolated ventilated caging system (Isocages from Techniplast, West Chester, Pennsylvania, USA, 2 mice per cage) to protect them from environmental contamination and in a controlled environment (12 hours day/night cycle, lights off at 7:00PM, 21°C±2°C, 42°C±13% of humidity). Mice had ad libitum access to autoclaved food and autoclaved drinking water and were between 8 and 10 weeks old at the start of the experiment. All procedures were approved by the animal experimentation committee of the Institut Pasteur and authorized by the French Ministry of Research (DAP n°220104)

#### CRISRPi screen *in vivo*

CRISPRi screens followed the experimental workflow described below. For each condition (strain or diet), 2 cages of 2 mice (4 mice in total) were used.

### In vitro pre-induction

One day prior to mouse gavage, 1 mL of glycerol stock of the CRISPRi library was thawed and used to inoculate 9 mL of LB medium supplemented with Cm. The culture was grown overnight at 37°C. The following day, 1 mL of the overnight culture was collected for plasmid extraction using a Miniprep kit (Macherey-Nagel). This sample, referred to as T_0_, represented the initial composition of the library prior to dCas9 pre-induction. The rest of overnight culture was diluted to an ODCCC of 0.2 in fresh LB supplemented with DAPG and incubated for 3 hours at 37°C to induce dCas9 expression before gavage.

### Mouse gavage

After 3 hours of *in vitro* dCas9 pre-induction, 1mL of the bacterial culture was kept for plasmid extraction. This sample, referred to as T_0_induced_, served as the reference time point for subsequent CRISPRi analysis. The remaining 9mL were washed once with 9 mL of sterile PBS and resuspended in 900 µL of PBS. Each mouse was gavaged with 200 µL of a preparation consisting of: 100 µL of concentrated bacteria in PBS (equivalent to ∼1 mL of overnight culture, ∼10C bacteria), mixed with 100 µL of a sucrose-bicarbonate buffer (200 mg/mL sucrose, 26 mg/mL NaHCOC in water). DAPG was added to the drinking water (provided ad libitum) at a final concentration of 1.2 mM and maintained throughout the experiment.

### Plasmid extraction from feces or gut content

Feces were collected at multiple time points from 6 hours up to 9 days post-gavage. At the end of the experiment, mice were sacrificed, and the gut was collected and divided into three regions: the colon, the cecum, and the small intestine. The contents of each region were collected separately.

Fecal pellets or gut content were weighed and rehydrated in 4 mL of PBS for 15 minutes. Samples were homogenized by vortexing and manually crushed using an inoculation loop. After homogenization, samples were allowed to decant for 15 minutes to separate large particles. The supernatant, containing the resuspended bacteria, was carefully transferred to a new tube. Serial dilutions of the supernatant in PBS were plated on Drigalski+Strep plates to quantify total *E. coli*, and Drigalski+Strep+Cm plates to quantify *E. coli* carrying the CRISPRi plasmid. Remaining bacterial suspensions were centrifuged at 4,000 g for 15 minutes, and the pellet was used for plasmid miniprep (NucleoSpin Plasmid kit, Macherey-Nagel). To maximize DNA recovery, three consecutive elutions with 30 µL of water were performed in the final step. ***Diets.*** For dietary experiments, the regular chow diet normally used in the animal facility was replaced with one of the following purified diets from Research Diets Inc., starting 3 days prior to *E. coli* gavage and maintained *ad libitum* throughout the experiment : high fat diet (Research Diets D12492i : Rodent Diet With 60 kcal% Fat), high fiber diet (Research Diets D13081108 : Rodent Diet With 10 kcal% Fat With No Cellulose and 200g Inulin per 4057 kcals) and standard diet (Research Diets D12450Ji : Rodent Diet With 10 kcal% Fat) (Table S2) ***DSS inflammation model.*** To induce colitis, 1.5% DSS (dextran sulfate sodium, 36,000-50000 Da, MP Biomedicals) was added to the drinking water starting 7 days before *E. coli* (MG655 or LF82) gavage. On the day of gavage, DSS-containing water was replaced with DAPG-containing water and maintained until the end of the experiment. Feces were collected daily throughout the experiment to monitor inflammation. To monitor inflammation, the presence of blood in stool, stool consistency, and mouse weight loss were analysed daily. One mouse colonized with the LF82 strain died before the end of the experiment.

#### CRISPRi screen *in vitro*

For each CRISPRi library, overnight cultures were initiated by inoculating 500 µL of glycerol stock into 5 mL of LB supplemented with Cm. After overnight growth at 37°C, cultures were washed once with LB to remove residual antibiotics. For the experimental condition, 15 µL of washed culture was transferred into 1,485 µL of LB supplemented with 50 µM DAPG in duplicate (biological replicates). The remaining overnight culture was collected, and plasmid DNA was extracted by miniprep (Macherey-Nagel) to serve as the T_0_ reference sample, representing the initial composition of the library. After three successive passages (1:100 dilution every 3.5 hours), corresponding to approximately 20 bacterial generations, plasmid DNA was extracted again to obtain the final time point (T_final_) composition of the library.

### Illumina library preparation and sequencing

Illumina library preparation and sequencing was performed as previously described^9^. A first PCR was performed on half of the miniprep product form CRISPRi screen (to avoid any bottleneck) using the PHUSION polymerase (Thermo Fisher) with LC863 and primers in the *PCR1_fwd* list (Table S1). This first PCR amplifies the sgRNAs region of the plasmid adding the 2 Unique Molecular Identifier (UMI) and a first index (PCR : 95 °C for 5 min; 2 cycles of 98C°C for 30 s, 60 °C for 90 s and 72 °C for 30 s; 72 °C for 5 min). The PCR product was digested by ExoI enzyme for 1h at 37°C, then cleaned up using AMPure XP beads (Beckman Coulter). Elution was done in 30µL, and 10µL of it was used for the second PCR done with LC415 and primers in the *PCR2_fwd* list in Table S1, to add the second index and the P5 and P7 Illumina adapters (PCR : 95 °C for 5 min; 12, 20 or 25 cycles of 98 °C for 30Cs, 60C°C for 90 s and 72 °C for 30 s; 72 °C for 5 min). For samples from bacterial culture (T_0_ or *in vitro* screen) 12 cycles were performed for the second PCR. For the *in vivo* screen, starting with less material from the feces or gut content, either 20 or 25 cycles was performed, eventually choosing the minimal number of cycles that allowed to get at least 0.2ng/µL of the 354 expected band. All PCR_2 samples were quantified using a TapeStation system (Agilent), normalized at 0.2ng/µL, and then pooled to perform a unique gel extraction of all the samples at once. Three elutions were made in 30µL without heating, to increase yield. The final illumina library was quantified by qPCR with NEBNext^®^ Library Quant Kit for Illumina^®^. 650pM of the library was sequenced using custom primers. LC610 and LC499 were used to read the indexes and the first UMI (*Index 1* and *Index 2* with 8 cycles). AM323 was used to read the second UMI (*Read 2* with 6 cycles). LC609 was used to read the sgRNA (*Read1* with 21 cycles). To avoid low diversity issues, a custom protocol was used to introduce 2 first dark cycles before additional 20 normal sequencing cycles to *Read 1* which reads the guide RNA sequence. The sequencing run was done on a NextSeq2000 sequencer at the Biomics platform of Pasteur Institute.

### Illumina sequencing data analysis

For the experiment comparing the three strains and the DSS-induced colitis model, the final time point for each mouse (D9 and D15, respectively) was used for the analysis presented in the figures. In the diet experiment, stronger CRISPRi signals (reflected by the depletion of sgRNAs targeting essential genes) were observed at earlier time points. Therefore, samples from D8 were used, except for the second cage in the high-fiber diet condition, where a bottleneck likely occurred during sample handling. In this case, samples from the final time point (D12), which showed a comparable signal strength to D5 in other cages, were used instead.

Signals also likely due to bottleneck were observed in 1 out of the 4 mice gavage with the CFT073 library, therefore this mouse was excluded from the analysis.

## Data Processing and Normalization

Raw BCL data generated by the Illumina sequencer were converted to FASTQ format using the bcl2fastq tool (version 2.20.0). A blank sample sheet containing a dummy index was used to prevent automatic demultiplexing, allowing all reads to be pooled into the Undetermined.fastq files. Index sequences were then used to manually demultiplex the sequencing data into individual samples using a custom Python script. For each sample, the script generated a CSV file containing the number of unique molecular identifiers (UMIs) and total read counts for each guide in the library. Guides with fewer than 20 UMIs at the initial time point were excluded from downstream analysis. A first analysis was done using the MAGeCK (version 0.5.9.5) pipeline to identify genes significantly essential or deleterious in each condition.Then, log2FC for each gene was calculated as follows to compare results across different conditions.

## Calculation of Log2 Fold Change (Log2FC)

To correct for sequencing depth and technical biases, the number of UMIs per guide was normalized to the total number of UMIs per sample using the DESeq2 package. For each guide, the log2 fold change (Log2FC) was calculated as follows:

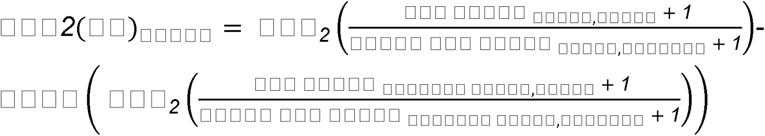

The Log2FC values were then centered on the mean Log2FC of the control guides as reference for a log2FC of 0 as they should not have an effect of bacterial growth. Then, the median Log2FC was calculated for each gene. Genes with less than two associated guides were excluded from further analysis. The final Log2FC dataset is available in the supplementary material.

### Identification of genes with significantly different signals across conditions

The overall strength of the depletion signal can be different in between experiments, likely due to the fact that E. coli does not perform the same number of generations in all conditions. To take this effect into account, for each comparison we first fitted a line to the median log2FC values of guides targeting each gene. We then tested the null hypothesis that the distance between guides targeting a gene and the fitted line was zero using one sample Student’s t-test. An FDR correction was then applied for multiple testing.

### Prophage induction quantification

Frozen fecal samples were thawed on ice for 10 min and diluted 40-fold in cold PBS. The suspension was incubated on ice for 5 min and centrifuged at 5000 × g for 10 min at 4°C. The pellet was used for genomic extraction using a method that was previously described^48^ and based on the Godon DNA extraction method^49^. The supernatant was filtered through a 0.22 μm filter and used for virome extraction.

#### Virome Extraction

For phage quantification by qPCR, virions were precipitated by adding polyethylene glycol (PEG) 8000 (10%) and NaCl (1 M), followed by overnight incubation at 4°C. Phage particles were collected by centrifugation at 5251 × g for 1 h at 4°C using a swinging rotor and resuspended in 100 μl of SM buffer (100 mM NaCl, 8CmM MgSO4, 50 mM Tris pH 7.5 (Sigma)). The samples were treated with 10 U of Turbo DNase (Ambion) for 1 h at 37°C in Turbo DNase buffer. The enzyme was inactivated, and capsids were disrupted by incubation at 95°C for 30 min in 0.2 mL PCR tubes.

#### Phage Quantification by qPCR

##### Template Preparation

For each sample, two dilutions of phage DNA extracted from feces (1:50 and 1:100) were prepared. 6 μl of each dilution was used for qPCR. 6 μl of water was used as negative control. Standards were prepared with LF82 genomic DNA (extracted using the Wizard DNA extraction kit (Promega) and quantified using a Qubit fluorometer (Invitrogen)). The DNA concentration was adjusted to obtain a solution containing 5 × 10^5^ genome copies. Serial 1:10 dilutions were performed four times to generate a standard curve ranging from 5 × 10^1^ to 5 × 10^5^ copies.

##### qPCR Reactions

5 qPCR reactions were conducted for 5 targets: Gally, Tritos, Perceval, Cartapus and LF82 genomic DNA (for genomic contamination control). The 5 pairs of primer for each qPCR run are in Table S1. Each qPCR run included one negative control (water), five standard dilutions (LF82 genome), and two dilutions per sample replicates (phage DNA extraction). The qPCR reaction mix (total volume of 20 μl) was prepared with 10 μl Luna qPCR mix (Luna® Universal qPCR Master Mix, NEB), 0.5 μl of each primer (250 nM), 6µL of DNA template or water, 3µL of nuclease-free water. The qPCR reaction was then incubated for 1min at 95°C and for 40 cycles of 15sec at 95°C and 30sec at 60°C. All reactions were performed in triplicate to ensure reproducibility. Tritos, Perceval, and Cartapus were not detected in sufficient amounts by qPCR, likely due to their low expression levels and the limited amount of material available in the fecal samples.The same qPCR reaction was performed on DNA extracted from the bacterial pellet of the same fecal samples, using LF82 genome specific primers, to estimate the absolute number of LF82 cells. The number of phage particles was then normalized to the number of LF82 clones in each feces sample.

### 16S analysis

Total DNA was extracted from feces using a method that was previously described^48^, which is based on the Godon DNA extraction method^49^. DNA concentration and quality were measured using a NanoDrop 2000 spectrophotometer (Thermo Fisher Scientific). Samples were sent to the Genoscreen platform (Lille, France) for the 16S rRNA sequencing on an Illumina MiSeq (Illumina, San Diego, CA, USA). Sequencing data were processed and analyzed using QIIME 2 (v. metagenome-2024.5) and its DADA2 pipeline. Taxonomic classification was performed using a custom classifier trained specifically on the 16S rRNA gene sequences of the 13 bacterial strains from the OligoMM12 consortium plus *Escherichia coli*.

## Acknowledgements

We thank all the members of the animal core facility (Institut Pasteur) for their assistance in mice care, Marie-Agnès Petit (INRAe, Jouy-en-Josas, France) for helpful discussions and advices, Andrew Gewirtz (Georgia State University, USA) for providing diets, Genoscreen (Lille, France), and Prebiomics (Trento, Italy) for assistance in sequencing, all the members of the Bikard and Seksik-Sokol team for technical and scientific discussions. This study was funded by European Research Council [101044479] to D.B., Agence Nationale de la Recherche [ANR-10-LABX-62-IBEID] to D.B., Fondation pour la Recherche Médicale [FDT202404018404] to A.M., Swiss National Science Foundation NCCR AntiResist (51NF40_180541) to M.O. and C.D.

## Author contributions

A.M., N.R. and D.B. conceived and designed the study. M.O. constructed the CFT073 guide RNA library. A.M. performed all the experiments with input from N.R. for mice experiments. A.M. analysed, interpreted the results and drafted the manuscript with input from all co-authors. E.T. provided technical help. B.C., C.D. and H.S. provided materials and intellectual input. All authors read and approved the final manuscript.

## Data and code availability

The CRISPRi raw sequencing data were deposited on SRA (Accession : PRJNA1293909). Scripts and notebooks used in data analysis are available at https://gitlab.pasteur.fr/dbikard/in-vivo-crispri.

