## Supplementary material for "Decoding *E. coli*’s Gut Survival Strategies: A CRISPRi Approach Across Diets, Inflammatory Environment and Strains"

### **Supplementary Text**

***Establishment of a CRISPRi screening strategy for E. coli in the mouse gut***

We first sought to identify experimental conditions suitable for CRISPRi screening in the animal gut. We assessed the ability of two different expression systems to induce dCas9 in the mouse gut environment when giving the inducer in the drinking water. In the first system, dCas9 was chromosomally integrated under the control of the anhydrotetracycline (aTc)-inducible pTet promoter. Two strains, ACE1^1^ and LC-E18^2^, were tested; they differ mainly in the ribosome binding site (RBS) used for dCas9 expression. LC-E18 harbors a strong RBS, resulting in lower dCas9 expression, whereas ACE1 contains a weaker RBS, leading to higher expression levels. Mice received 30 μg/mL of aTc in their drinking water to induce dCas9. In the second system, dCas9 was expressed from the plasmid pFR56 under the control of the DAPG-inducible pPhlF promoter. This plasmid also carried the sgRNA. Various DAPG concentrations (25 µM, 250 µM, and 2.5 mM) were tested via drinking water. However, in both systems and across all inducer concentrations, CRISPRi screens failed to show the expected depletion of sgRNAs targeting essential genes, suggesting that dCas9 expression in the gut environment remained insufficient (Fig. S1A–D).

To overcome this limitation, we introduced a 2H in vitro pre-induction step prior to oral gavage, ensuring the bacteria were expressing dCas9 at the time of administration. The phenotypic effect of dCas9 induction on bacterial fitness is not immediately apparent, but rather takes several generations to manifest. This delay occurs because dCas9 must be produced to sufficient levels, bind its target on the chromosome, and then existing transcripts and proteins must be degraded or diluted before an effect is observed. We thus reasoned that this short pre-induction would have a minimal effect on the overall signal obtained in the screens. Supporting this, comparing the CRISPRi library composition before and after pre-induction revealed no significant depletion of any guide RNAs. The inducer (DAPG) was then maintained in the mice's drinking water throughout the experiment. This protocol resulted in a clear depletion of guides targeting essential genes and showed a strong correlation between biological replicates, demonstrating a successful in vivo CRISPRi screen (Fig. S1EF).

***Deleterious genes reveal the misadaptation of MG1655 to the gut environment***

In the mouse gut, MG1655 strongly benefits from dCas9 silencing of the transcriptional activator of the maltose transporters (*malT*), and of the maltose transporters themselves (*malEFG* and *malK-lamB* operons). This aligns with previous findings that MG1655 rapidly accumulates mutations in these genes when colonizing the gut^3^. Interestingly genes involved in maltose utilisation such as *malQ* and *glk* are however important for E. coli’s fitness in the gut. A possible explanation for this discrepancy is that the shutdown of porins might help protect against osmolarity conditions found in the gut, while the ability to metabolize maltose remains beneficial. Similarly a strong signal is also observed for *gntT* (gluconate transporter), with the opposite effect observed for *gntR*, the repressor of *gntT*.

Among genes whose expression is deleterious to MG1655 across diets, we also find several global regulators. This includes *hns*, which was described to play a role in the regulation of carbon and nitrogen metabolism as well as response to multiple stresses^4^^,^^5^. Another strong signal comes from *sspA* which was described to play a role in the adaptation to acid stresses^6^. SspA is encoded in an operon with *sspB* for which a strong signal is also observed. The SspB protein promotes the degradation of specific proteins by ClpXP, and in particular of the anti-sigma E factor RseA, which is involved in the response to multiple stresses including hyperosmotic shock^7^. Another global regulator found to be deleterious to MG1655 in the mouse gut is the *rhoL-rho* operon. Rho is involved in transcription termination and is essential to E. coli during in vitro growth, it is thus surprising to find it as deleterious in vivo. Previous work on *rho* mutants showed an alteration of the outer membrane and increased adhesion of MG1655, which could favor gut colonization, but an increased susceptibility to bile acids, which should have the opposite effect^8^. While more work will be necessary to understand the mechanism underlying the effects of these regulators, our results show how MG1655 largely fails to adjust its gene expression profile to in vivo growth, where it competes for nutrients with other bacteria and encounters various stresses. As a result, silencing key global regulators with dCas9 can facilitate growth in this environment.

Strong signals observed for other genes involved in metabolism are harder to explain and thus not discussed in the main text. This includes the deleterious effect of the *pykA* pyruvate kinase, and of the *plaR* repressor of the rare sugar L-lyxose catabolism in the standard diet.

***Sorbitol cross-feeding in the high-fiber and standard diets***

Silencing *srlR* in standard and high-fiber diets, thereby activating the sorbitol utilisation operon is highly beneficial in these environments. Since sorbitol is not present in these diets, we can hypothesize that it is produced from glucose by gut bacteria through the polyol or glucose-fructose oxidoreductase (GFOR) pathways. Supporting this, oxidoreductases from these pathways can be found in several members of the OligoMM12 community. This observation on *srlR* is also consistent with a previous report of srlR inactivation through mutations during E. coli adaptation to the mouse gut environment^9^.

***Different stress adaptation pathways are needed to adapt to environmental changes in the high fat-diet***

In addition to differences in nutrient availability, several lines of evidence suggest that the high-fat diet induces greater environmental changes in the gut, impacting E. coli stress response more than the other diets. First, the high-fat diet environment appears to be lower in oxygen. The *fnr* regulator, which activates anaerobic metabolism genes and silences aerobic metabolism genes is more important to MG1655’s fitness under this diet, with the small regulatory RNA *fnrS* showing a weaker but consistent signal. The activity of *fnr* requires the presence of an intact Fe-S cluster. Consistently, the genes involved in the ISC Fe-S cluster generation machinery (*iscRSUA-hscBA-fdx-iscX*) are more important for fitness in this environment than in the other two diets. Similarly the ArcAB two component system which is active under anaerobic conditions and regulates the expression of several operons involved in respiratory and fermentative metabolism plays an important role in the high-fat diet.
Another specificity of the high-fat diet is the importance of NhaA, which imports protons and exports sodium ions, helping maintain internal pH balance when the environment becomes alkaline. A *nhaA* mutant was also found to be more susceptible to sodium dodecyl sulfate, suggesting an impaired envelope integrity. High-fat diets stimulate increased bile acid secretion into the intestine which could both create a more alkaline environment in the gut and kill E. coli cells with a destabilized envelope.
E. coli further appears to adapt to these environmental changes through genes involved in stress response (*rpoH, hflXKC, lexA*), motility regulation (*lrhA*), and global transcriptional regulation (*ihfAB*), which become important under the high fat diet only.

***Cost of flagellum in MG1655 in the chow diet***

We observe that flagellar genes are deleterious for MG1655 in the gut of mice fed with a chow diet. E. coli was previously found to rapidly mutate the *flhDC* genes during adaptation to the mouse gut environment^3,10^. Those mutations have been shown to increase carbon and energy metabolism^11^, which could explain the observed fitness advantage of silencing flagellar genes. Interestingly, this signal was not observed when mice were fed purified diets, suggesting that diets impact the properties of the environment, either increasing the level of expression of the flagella to maladaptive levels, or making motility less beneficial in the chow diet. Conversely, the neutral signal for those genes in LF82 and CFT073 could indicate that they regulate flagella expression in an appropriate manner, shutting down their expression when not needed, or that these strains benefit more from motility than MG1655, compensating the cost of flagella expression. Indeed, the flagellum is a known virulence factor in LF82 and CFT073, and facilitates gut colonization of pathogens in general^12^^,^^13^.

***Expression of galactitol operon in the gut***

The role of *gatR*, which is non-functional in MG1655 and appears to be strongly deleterious for CFT073 and LF82 in vivo, has been frequently discussed in the literature. We both read that MG1655 can rapidly mutate gat genes leading to the emergence of a gat-negative phenotype^14^, and that gatR inactivation in a gatR-restored MG1655 rapidly re-emerge in vivo^15^. These opposite observations do suggest that the advantage of gat utilization ability depends on the environment : *gat* metabolism can be deleterious, possibly due to the energy cost of constitutive expression or the toxicity of galactitol metabolism intermediates, but it can represent a fitness advantage in environments where galactitol is sufficiently abundant.

This leads to the following overall model: in the gut of OMM12 mouse eating a chow diet, galactitol, which is a product of the host galactose metabolism that can vary depending on host genetic, diet or microbiota composition, is highly abundant, but LF82 and CFT073 fail to derepress the gat operon, even though they would benefit from metabolizing galactitol.

***Deleterious metabolic regulator in LF82 :***

Similarly to *gatR*, two other regulators of metabolic operons are deleterious to LF82 only: *dgoR* and *galR*, the repressors of operons involved in galactonate and galactose metabolism respectively. Consistently, previous studies have shown that mutations in *dgoR* and an enhanced ability to utilize galactose are selected in E. coli during evolution in the gut of mice^16^. Curiously, we also found several sugar transporters (*fruA, gntT,* and *manZ*) to be deleterious to LF82, which is reminiscent of the deleterious effect of maltose transport in MG1655, even if the signal is weaker. This could indicate that LF82 suffers from the toxic accumulation of unused carbohydrates in the gut environment.

***Chemical stress in the gut environment***

Another gene which is deleterious across strains is *emrR*, the repressor of the emrAB operon encoding an efflux pump involved in multidrug resistance^17^, suggesting that E. coli benefits from increased expression of this transporter. We also observed the same signal for ygaH, located directly upstream of the promoter of emrR and which encodes a membrane transporter for L-valine involved in susceptibility to environmental stress signals (phages, antibiotics, and salt)^18–20^. These observations further highlight the importance of chemical stress resistance in the gut environment.

***Differences in Respiration among the 3 strains***

We observe significant differences on genes involved in ECT across the three strains, suggesting they rely on different types of respiration. The most notable variation occurred for the dehydrogenases that catalyze the first step of respiration. Genes of the *nuo* operon, which form the NADH dehydrogenase I (NDH-I) complex, were substantially more critical for the fitness of MG1655 than for LF82 and CFT073,. In contrast, the *ndh* gene, which encodes NADH dehydrogenase II (NDH-II), appears deleterious in CFT073 but neutral in LF82 and MG1655. NDH-II is the predominant NADH dehydrogenase during aerobic growth when rapid NADH oxidation is prioritized and energy is not strictly limiting, while NDH-I functions under both aerobic and anaerobic conditions^21^. Crucially, NDH-I translocates protons across the cytoplasmic membrane, thereby directly contributing to the proton motive force essential for ATP synthesis, making it preferable under energy-limited conditions^22,23^. The reliance of MG1655 on NDH-I likely indicates that it operates under significant energy stress, consistent with the high cost observed for the expression of the flagellar genes. Other electron donors associated with anaerobic respiration, such as hydrogenase I and II (encoded by *hyb* and *hyo* genes, respectively), appear to be more important for CFT073. This observation, together with the deleterious nature of *ndh* for this strain, suggest it may inhabit a more anaerobic niche than MG1655 and LF82. The importance of hydrogenase also suggests the presence of hydrogen in the gut, likely produced by fermentation of the gut bacteria.

Signals from genes encoding terminal reductases and oxidases corroborate these observations. Under aerobic or microaerobic conditions, cytochromes bo and bd use oxygen as the terminal electron acceptor. Cytochrome bo is expressed in high-oxygen environments, while cytochrome bd-I, which has a higher affinity for oxygen, is expressed under oxygen-limited conditions^24^. The *cyo* genes, encoding for cytochrome bo, is more important for MG1655 than for LF82 and CFT073. In contrast, cyd genes, coding for the cytochrome bd-I complex, seem to be more strongly essential for LF82 and CFT073. This suggests that MG1655 relies more on aerobic respiration.

***The envelope modulation and stress response becomes important during inflammation***

An increased importance of genes involved in envelope stress response and LPS biosynthesis is observed in DSS condition. The opposite signal is observed for *ompA*, which suggests that lowering the expression of *ompA* in conditions of membrane stress is beneficial, which could occur if misfolded OmpA accumulates in the membrane. Paradoxically, silencing the *mlaFEDCB* operon, which normally maintains outer membrane lipid asymmetry, also conferred a fitness advantage, suggesting its function may be energetically costly or counterproductive under inflammatory stress.

Another possible explanation for the increased importance of LPS biosynthesis genes in the inflamed gut is that modulation of LPS structure is a well-established strategy used by bacteria to evade immune detection^25^. In addition, several genes involved in resistance to human antimicrobial peptides are closely linked to LPS biosynthesis^26^. Their increased importance under inflammatory conditions may therefore reflect a role in modifying LPS to escape enhanced immune surveillance. Similarly, the increased importance of genes such as *yehD* and *tamA* suggests a potential enhancement of biofilm formation capacity, a strategy previously shown to protect bacteria from environmental stress and host immune responses^27^ ^28^.

***Importance of genes from accessory genome in CFT073***

While mobile genetic elements and prophages of LF82 are discussed in the main text, several interesting signals also emerged from the analysis of CFT073's accessory genome.

Prophage repressors were among the strongest signals in CFT073. As expected, silencing the repressors of prophages 1 and 2 had a strongly deleterious effect, consistent with the induction of lytic genes. Interestingly, guides targeting the repressor of the third CFT073 prophage were specifically depleted in vivo but not in vitro (Fig. S4B). This suggests that repression of the repressor alone is insufficient to trigger prophage induction under in vitro conditions, and that additional signals present in vivo are required to activate the lytic cycle.

Furthermore, several transposase genes from the CFT073 accessory genome appeared important for fitness in vivo but were neutral in vitro (Fig. S4B), suggesting a role for genome rearrangement in adaptation to the gut environment.

28.	Biofilm formation as a novel phenotypic feature of adherent-invasive Escherichia coli(AIEC) | BMC Microbiology | Full Text. https://bmcmicrobiol.biomedcentral.com/articles/10.1186/1471-2180-9-202.

**Supplementary Tables**

**Table 1 :** Primers used in the study

|  | **qPCR primers** |  |  |  |
| --- | --- | --- | --- | --- |
| AM356 | GGCTCAGCGCGTGGAA | LF82 ybtE |  |  |
| AM357 | CGGCCAGTGGTCCAGAAA | LF82 ybtE |  |  |
| AM358 | GACGCCATCGACATACAGG | LF82 ybtE |  |  |
| AM359 | GCCTTGCGTCATCTTCTCCA | Gally injection |  |  |
| AM360 | TCTGAGCAACGCTGTTAGGG | Gally injection |  |  |
| AM361 | GCATGGGGGCCTTCTGTAA | Perceval vebk |  |  |
| AM362 | GCCAGCGATTTCACTTATCCC | Perceval vebk |  |  |
| AM363 | CATCCCGGTGACCATGCC | Tritos minor tail |  |  |
| AM364 | ACGGGATTTGAACTGAACGGTA | Tritos minor tail |  |  |
| AM365 | TTGTCCAGCGGTTGTTTACCT | Cartapus tail |  |  |
| AM366 | CGGCACTGGATACACTGAAC | Cartapus tail |  |  |
|  | **Illumina library Preparation** |  |  |  |
|  | **Sequence**  Color code : Index Adapters | **Role** | **PCR** | **index** |
| LC606 | TTCCCTACACGACGCTCTTCCGATCTTAGANNNNGCACGCCCGTCGCTCAGTCCTAGGTATAATACTA | Index 1 for custom sample preparation | PCR1 Fwd | TAGA |
| LC607 | TTCCCTACACGACGCTCTTCCGATCTCTCTNNNNGCACGCCCGTCGCTCAGTCCTAGGTATAATACTA | Index 1 for custom sample preparation | PCR1 Fwd | CTCT |
| LC608 | TTCCCTACACGACGCTCTTCCGATCTATTCNNNNGCACGCCCGTCGCTCAGTCCTAGGTATAATACTA | Index 1 for custom sample preparation | PCR1 Fwd | ATTC |
| FR230 | TTCCCTACACGACGCTCTTCCGATCTTGCTNNNNGCACGCCCGTCGCTCAGTCCTAGGTATAATACTA | Index 1 for custom sample preparation | PCR1 Fwd | TGCT |
| FR231 | TTCCCTACACGACGCTCTTCCGATCTCTAANNNNGCACGCCCGTCGCTCAGTCCTAGGTATAATACTA | Index 1 for custom sample preparation | PCR1 Fwd | CTAA |
| FR232 | TTCCCTACACGACGCTCTTCCGATCTAGAGNNNNGCACGCCCGTCGCTCAGTCCTAGGTATAATACTA | Index 1 for custom sample preparation | PCR1 Fwd | AGAG |
| FR233 | TTCCCTACACGACGCTCTTCCGATCTACTGNNNNGCACGCCCGTCGCTCAGTCCTAGGTATAATACTA | Index 1 for custom sample preparation | PCR1 Fwd | ACTG |
| FR234 | TTCCCTACACGACGCTCTTCCGATCTCGTCNNNNGCACGCCCGTCGCTCAGTCCTAGGTATAATACTA | Index 1 for custom sample preparation | PCR1 Fwd | CGTC |
| LC863 | GTGACTGGAGTTCAGACGTGTGCTCTTCCGATCTNNNNNNAAAGGACCCGTAAAGTGATAATGAT | Reverse primer for sequencing PCR1 | PCR1 Rev |  |
| LC416 | CAAGCAGAAGACGGCATACGAGATCGAGTAATGTGACTGGAGTTCAGACG | Index 2 for custom sample preparation | PCR2 Fwd | CGAGTAAT |
| LC417 | CAAGCAGAAGACGGCATACGAGATTCTCCGGAGTGACTGGAGTTCAGACG | Index 2 for custom sample preparation | PCR2 Fwd | TCTCCGGA |
| LC418 | CAAGCAGAAGACGGCATACGAGATAATGAGCGGTGACTGGAGTTCAGACG | Index 2 for custom sample preparation | PCR2 Fwd | AATGAGCG |
| LC419 | CAAGCAGAAGACGGCATACGAGATGGAATCTCGTGACTGGAGTTCAGACG | Index 2 for custom sample preparation | PCR2 Fwd | GGAATCTC |
| LC420 | CAAGCAGAAGACGGCATACGAGATTTCTGAATGTGACTGGAGTTCAGACG | Index 2 for custom sample preparation | PCR2 Fwd | TTCTGAAT |
| LC421 | CAAGCAGAAGACGGCATACGAGATACGAATTCGTGACTGGAGTTCAGACG | Index 2 for custom sample preparation | PCR2 Fwd | ACGAATTC |
| LC422 | CAAGCAGAAGACGGCATACGAGATAGCTTCAGGTGACTGGAGTTCAGACG | Index 2 for custom sample preparation | PCR2 Fwd | AGCTTCAG |
| LC423 | CAAGCAGAAGACGGCATACGAGATGCGCATTAGTGACTGGAGTTCAGACG | Index 2 for custom sample preparation | PCR2 Fwd | GCGCATTA |
| LC1508 | CAAGCAGAAGACGGCATACGAGATCTGATTAAGTGACTGGAGTTCAGACG | Index 2 for custom sample preparation | PCR2 Fwd | CTGATTAA |
| LC1509 | CAAGCAGAAGACGGCATACGAGATTAAGCAGTGTGACTGGAGTTCAGACG | Index 2 for custom sample preparation | PCR2 Fwd | TAAGCAGT |
| LC1510 | CAAGCAGAAGACGGCATACGAGATGAGCTACGGTGACTGGAGTTCAGACG | Index 2 for custom sample preparation | PCR2 Fwd | GAGCTACG |
| LC1511 | CAAGCAGAAGACGGCATACGAGATAGAGAGGCGTGACTGGAGTTCAGACG | Index 2 for custom sample preparation | PCR2 Fwd | AGAGAGGC |
| LC1512 | CAAGCAGAAGACGGCATACGAGATCCTACGGCGTGACTGGAGTTCAGACG | Index 2 for custom sample preparation | PCR2 Fwd | CCTACGGC |
| LC1513 | CAAGCAGAAGACGGCATACGAGATCCTCGATGGTGACTGGAGTTCAGACG | Index 2 for custom sample preparation | PCR2 Fwd | CCTCGATG |
| LC1514 | CAAGCAGAAGACGGCATACGAGATCAGTCGAGGTGACTGGAGTTCAGACG | Index 2 for custom sample preparation | PCR2 Fwd | CAGTCGAG |
| LC1515 | CAAGCAGAAGACGGCATACGAGATTTAATCGCGTGACTGGAGTTCAGACG | Index 2 for custom sample preparation | PCR2 Fwd | TTAATCGC |
| LC1516 | CAAGCAGAAGACGGCATACGAGATGTCGTGATGTGACTGGAGTTCAGACG | Index 2 for custom sample preparation | PCR2 Fwd | GTCGTGAT |
| LC415 | AATGATACGGCGACCACCGAGATCTACACTCTTTCCCTACACGACGCT | Forward primer for sequencing PCR2 | PCR2 Rev |  |
| LC609 | GCACGCCCGTCGCTCAGTCCTAGGTATAATACTA | Custom read 1 primer for sequencing | oligos illumina sequencing |  |
| LC499 | GATCGGAAGAGCACACGTCTGAACTCCAGTCAC | Index primer 1 for sequencing | oligos illumina sequencing |  |
| LC610 | TATTATACCTAGGACTGAGCGACGGGCGTGC | Index primer 2 for sequencing | oligos illumina sequencing |  |
| AM323 | GTGACTGGAGTTCAGACGTGTGCTCTTCCGATCT | Custom read 2 primer for sequencing (= Illumina standard primer) | oligos illumina sequencing |  |
|  | **Cloning** |  |  |  |
| Sol70 | agctgtcacaggtctcaTAG | Amplify the CRISPRi library |  |  |
| Sol71 | gtagacgtagtcgcggtctc | Amplify the CRISPRi library |  |  |
| AM283 | G*T*CAGACGAACACGGCATACTTTACGCAGCGCGGAGTTCGGTTTTCTAGGAGTTGTAGTATATACACGAGTACATACGCCACGTTTTTGC | Introduce K43R mutation in rpsL with pKOBEG |  |  |

**Table 2 :** Diets composition

|  |  | **High Fiber** | | **Standard** | | **High-fat** | |
| --- | --- | --- | --- | --- | --- | --- | --- |
|  |  | g | % | g | % | g | % |
| Protein | Casein, Lactic, 30 Mesh | 200 | 17.32% | 200 | 18.96% | 200 | 25.84% |
| Protein | Cystine, L | 3 | 0.26% | 3 | 0.28% | 3 | 0.39% |
| Carbohydrate | Starch, Corn | 456.2 | 39.50% | 506.2 | 47.98% | 0 | 0.00% |
| Carbohydrate | Lodex 10 | 125 | 10.82% | 125 | 11.85% | 125 | 16.15% |
| Carbohydrate | Sucrose, Fine Granulated | 72.8 | 6.30% | 72.8 | 6.90% | 72.8 | 9.41% |
| Fiber | Solka Floc, FCC200 | 0 | 0.00% | 50 | 4.74% | 50 | 6.46% |
| Fiber | Raftiline HP | 200 | 17.32% | 0 | 0.00% | 0 | 0.00% |
| Fat | Soybean Oil, USP | 25 | 2.16% | 25 | 2.37% | 25 | 3.23% |
| Fat | Lard | 20 | 1.73% | 20 | 1.90% | 245 | 31.66% |
| Mineral | [S10026B](https://researchdiets.com/en/formulas/S10026B) | 50 | 4.33% | 50 | 4.74% | 50 | 6.46% |
| Vitamin | Choline Bitartrate | 2 | 0.17% | 2 | 0.19% | 2 | 0.26% |
| Vitamin | [V10001C](https://researchdiets.com/en/formulas/V10001C) | 1 | 0.09% | 1 | 0.09% | 1 | 0.13% |
| Dye | Dye, Blue FD&C #1, Alum. Lake 35-42% | 0.03 | 0.00% | 0.04 | 0.00% | 0.05 | 0.01% |
| Dye | Dye, Yellow FD&C #5, Alum. Lake 35-42% | 0.03 | 0.00% | 0.01 | 0.00% |  | 0.00% |
|  |  | 1155.06 | 1 | 1055.05 | 1 | 773.85 | 1 |

**Supplementary Figures :**

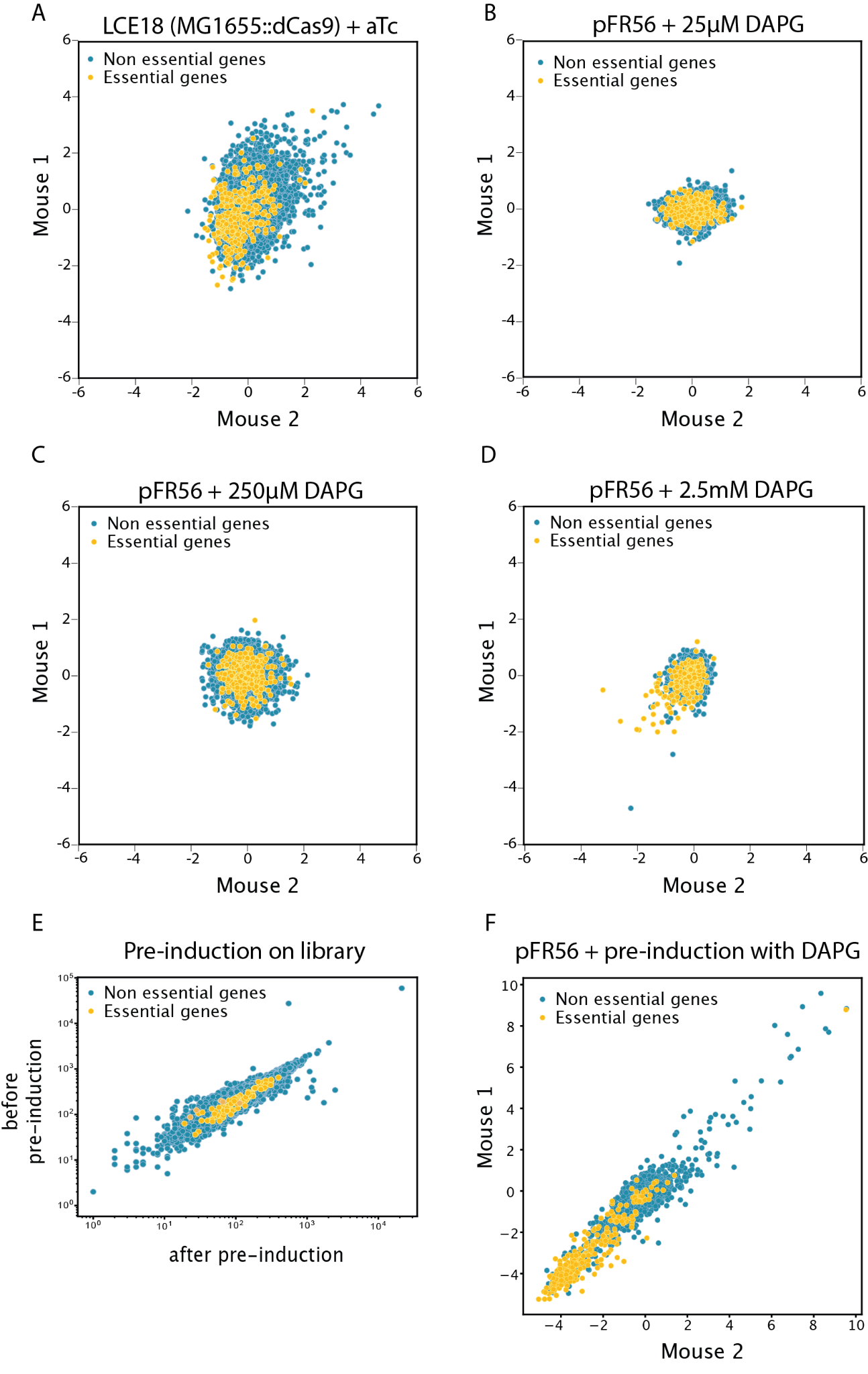
**Supplementary Figure 1. Optimization of *in vivo* expression of dCas9.**

Scatter plots of gene log₂ fold change (log₂FC) in MG1655 from one biological replicate (mouse 1) versus the other (mouse 2 from the same cage), assessing the setup’s ability to deplete guides targeting essential genes (in orange, expected to show low negative log₂FC values) and the overall reproducibility between replicates. (**A)** dCas9 expressed from the chromosome, induced with 30 ng/μL of anhydrotetracycline (ATc) in the mice drinking water. **(B-D)** dCas9 expressed from a plasmid under a DAPG-inducible promoter, tested with different DAPG concentrations: **(B)** 25µM, **(C)** 250µM, and **(D)** 2.5mM in the mice drinking water. **(E–F)** Setup with plasmid-encoded dCas9 and 2-hour *in vitro* pre-induction before gavage. **(E)** Scatter plot of guide abundance in the library before and after pre-induction, showing a strong correlation and confirming that pre-induction does not bias library composition. **(F)** Scatter plot of gene log₂FC in mouse 1 versus mouse 2, showing effective depletion of guides targeting essential genes (orange) and strong correlation between biological replicates.

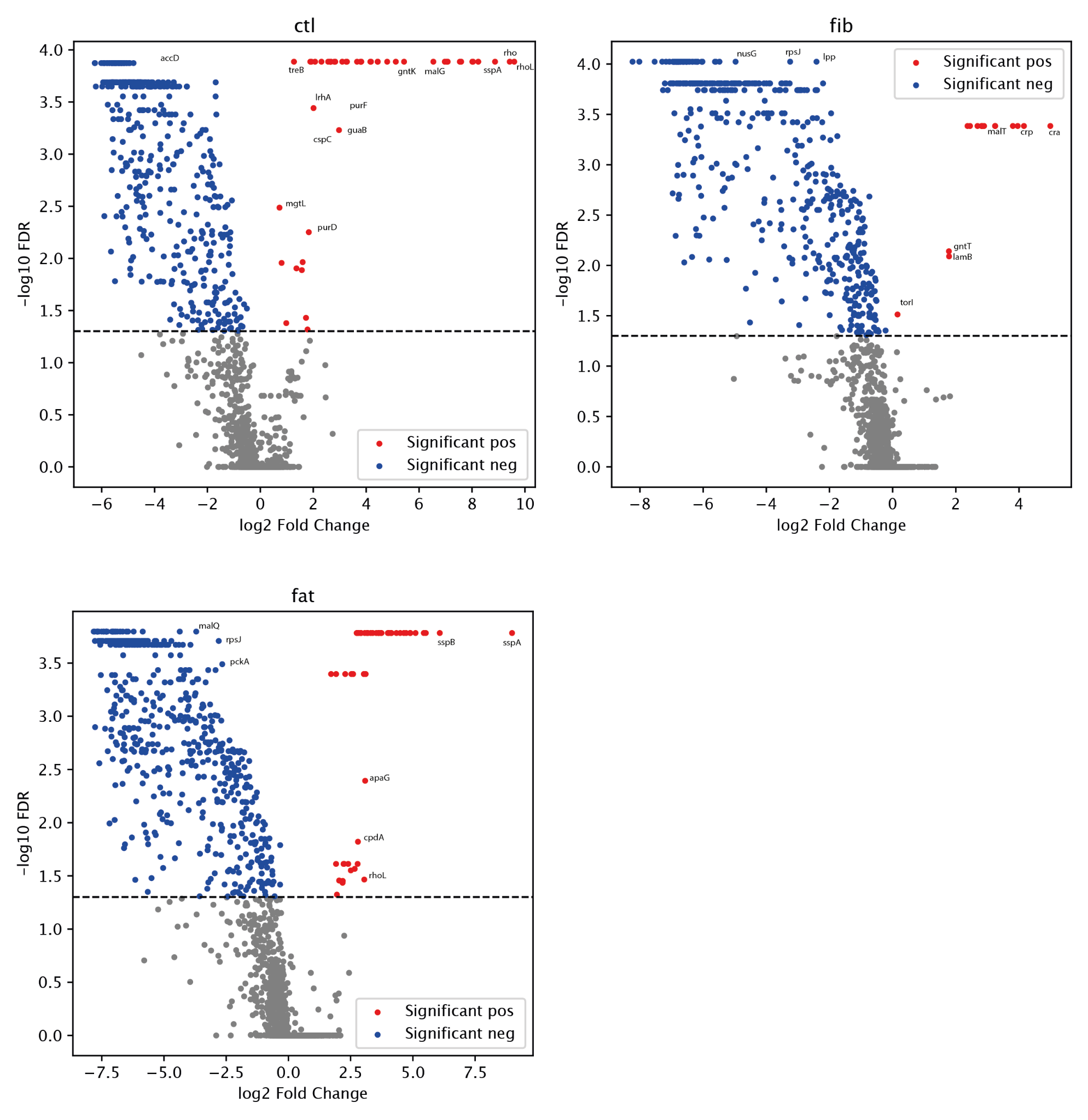

**Supplementary Figure 2. Volcano plot showing significantly essential and deleterious genes for MG1655 in each dietary conditions**

-log10(FDR) obtained for each gene with Mageck analysis in function of the gene log2FC. Red genes are significantly deleterious, blue genes are significantly essential.

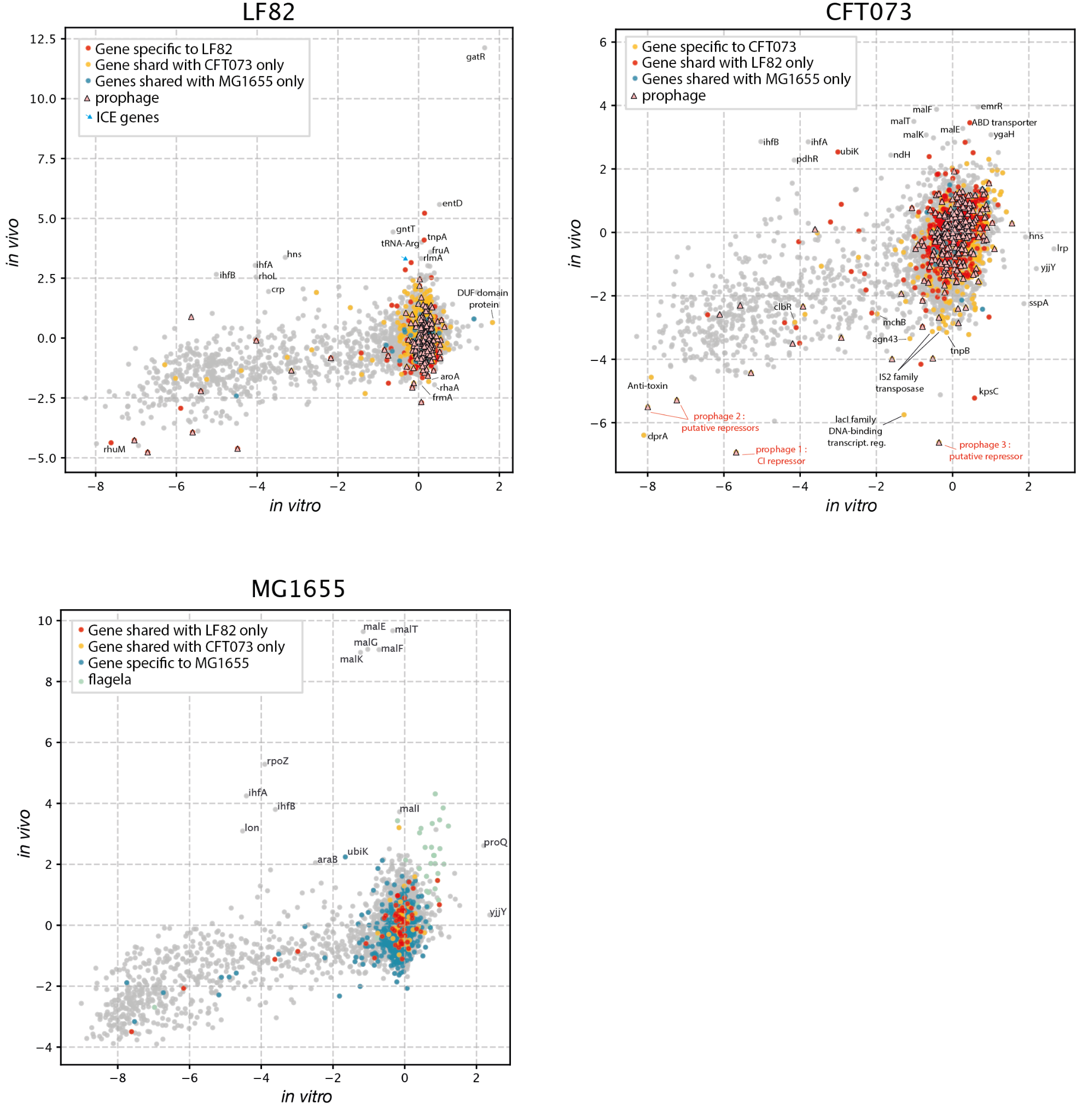

**Supplementary Figure 3. Comparison of gene essentiality in LF82 and CFT073 *E. coli* strains *in vitro* and in the gut.**

**(A–C)** Scatter plots of gene log₂ fold change (log₂FC) from *in vitro* screens in LB versus *in vivo* screens from feces of chow diet–fed mice without inflammation for LF82 **(A)**, CFT073 **(B)** and MG1655 **(C)**. Grey dots represent genes shared by all three strains, while colored dots indicate genes specific to the strain or shared with only one of the other two strains. Triangles represent genes located within prophage regions in (A) and (B).

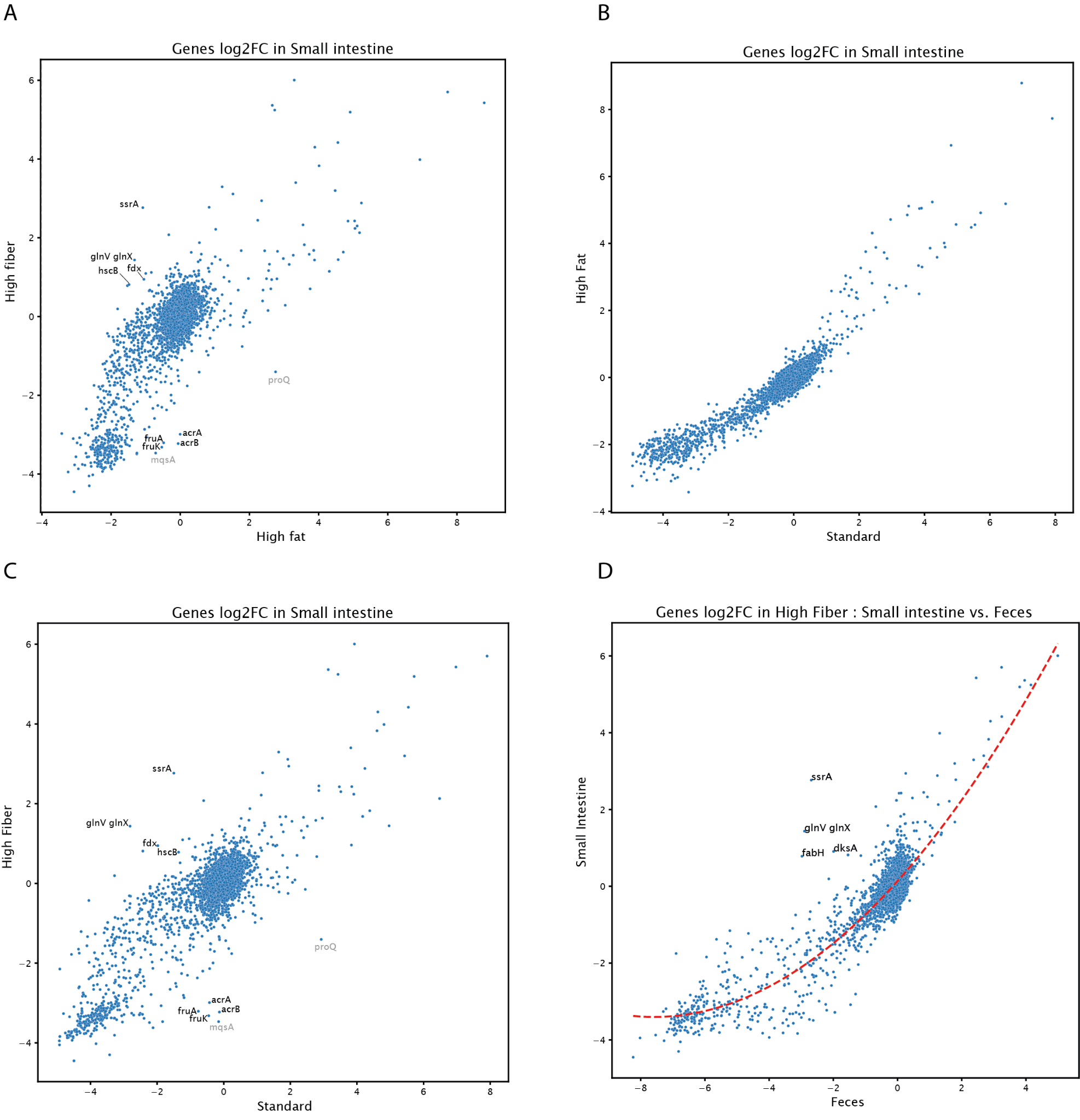

**Supplementary Figure 4. Gene essentiality of MG1655 in the small intestine.**
 **(A–C)** Pairwise comparisons of gene log₂ fold change (log₂FC) in the small intestine between the three dietary conditions. **(D)** Comparison of gene log₂FC between feces and the small intestine under the High Fiber diet condition.

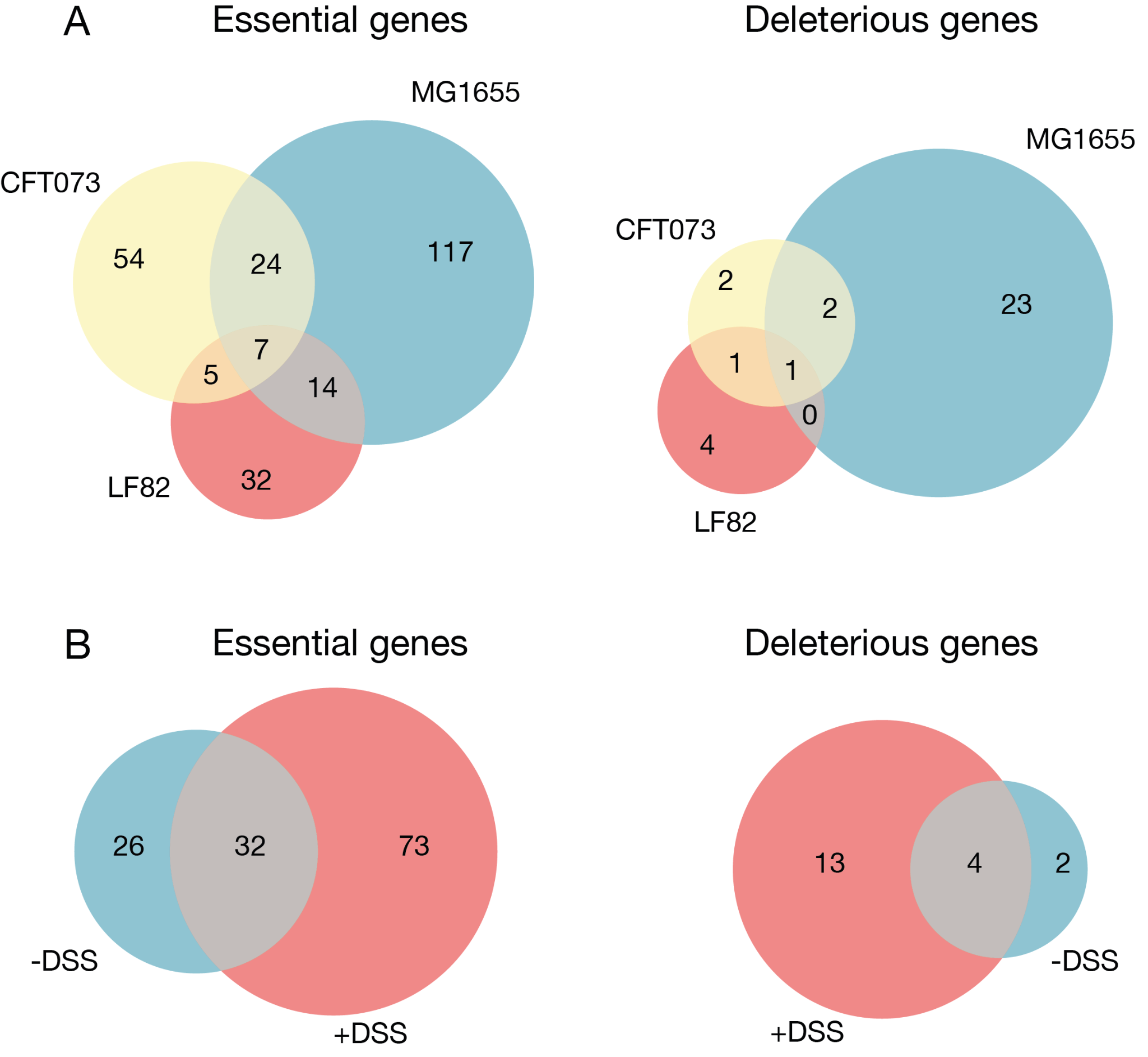

**Supplementary Figure 5. Number of significantly essential and deleterious genes in each conditions**
Number of genes with FDR < 0.05 in the MAGeCK analysis of positively selected genes (deleterious genes, right) or negatively selected genes (essential genes, left).

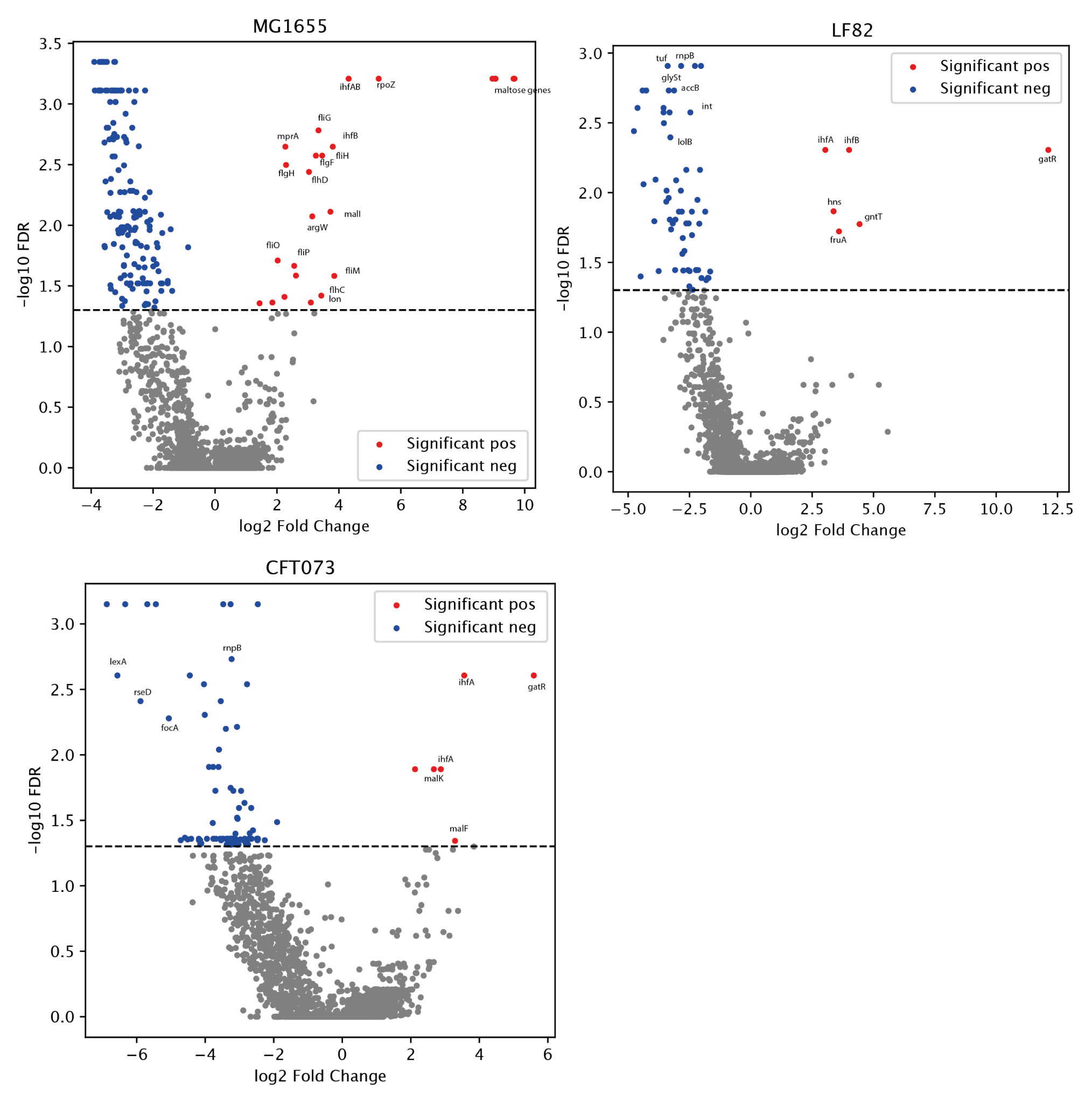

**Supplementary Figure 6. Volcano plot showing significantly essential and deleterious genes for each strain**

-log10(FDR) obtained for each gene with Mageck analysis in function of the gene log2FC. Red genes are significantly deleterious, blue genes are significantly essential.

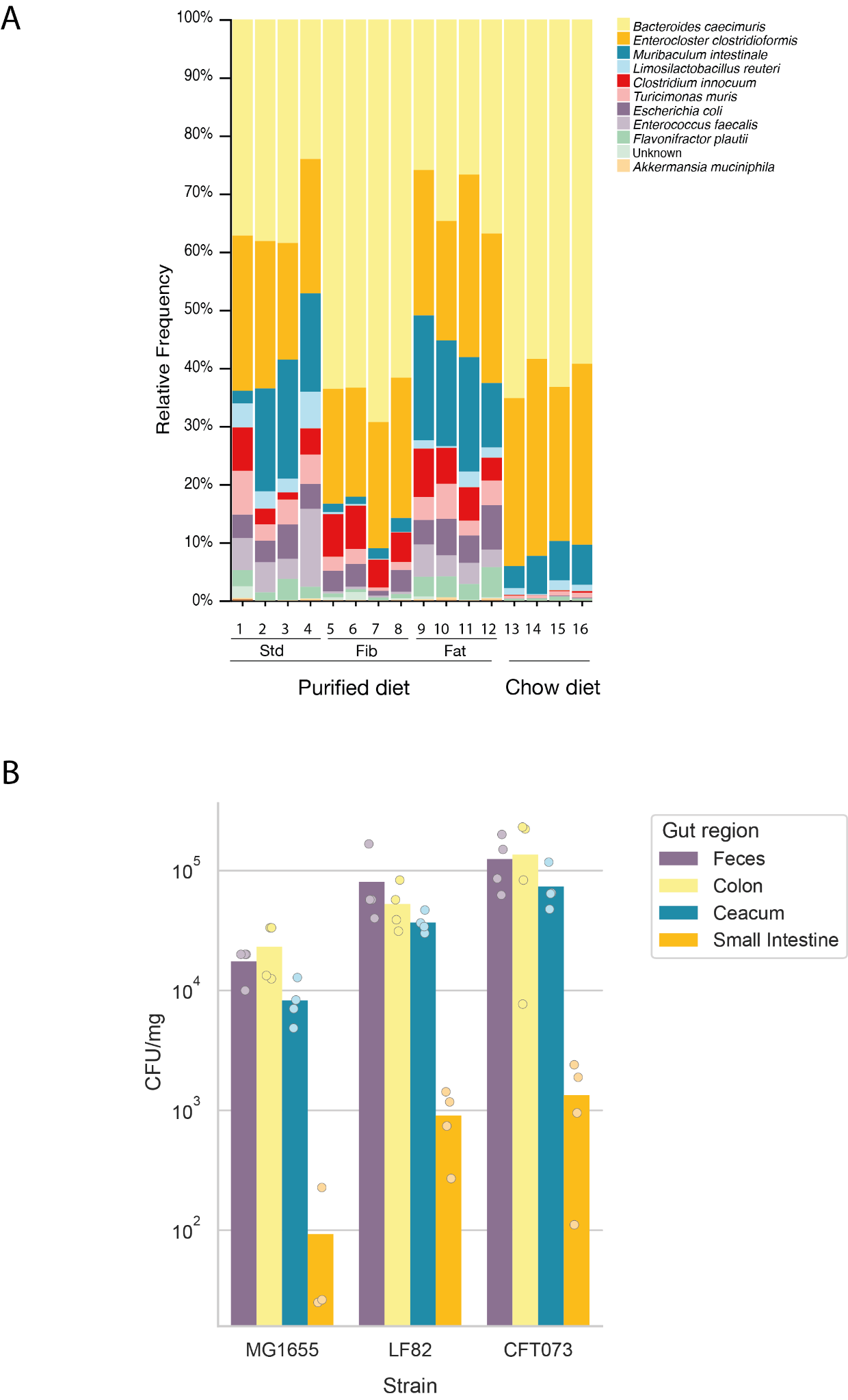

**Supplementary Figure 7. Colonization levels of MG1655, LF82, and CFT073 in different conditions.**

**(A)** Colonization levels of *E. coli* strains MG1655 across different dietary condition **(B)** Colonization levels of *E. coli* strains MG1655, LF82, and CFT073 in various gut regions of chow diet–fed mice without inflammation induction, expressed as CFU per mg of content.

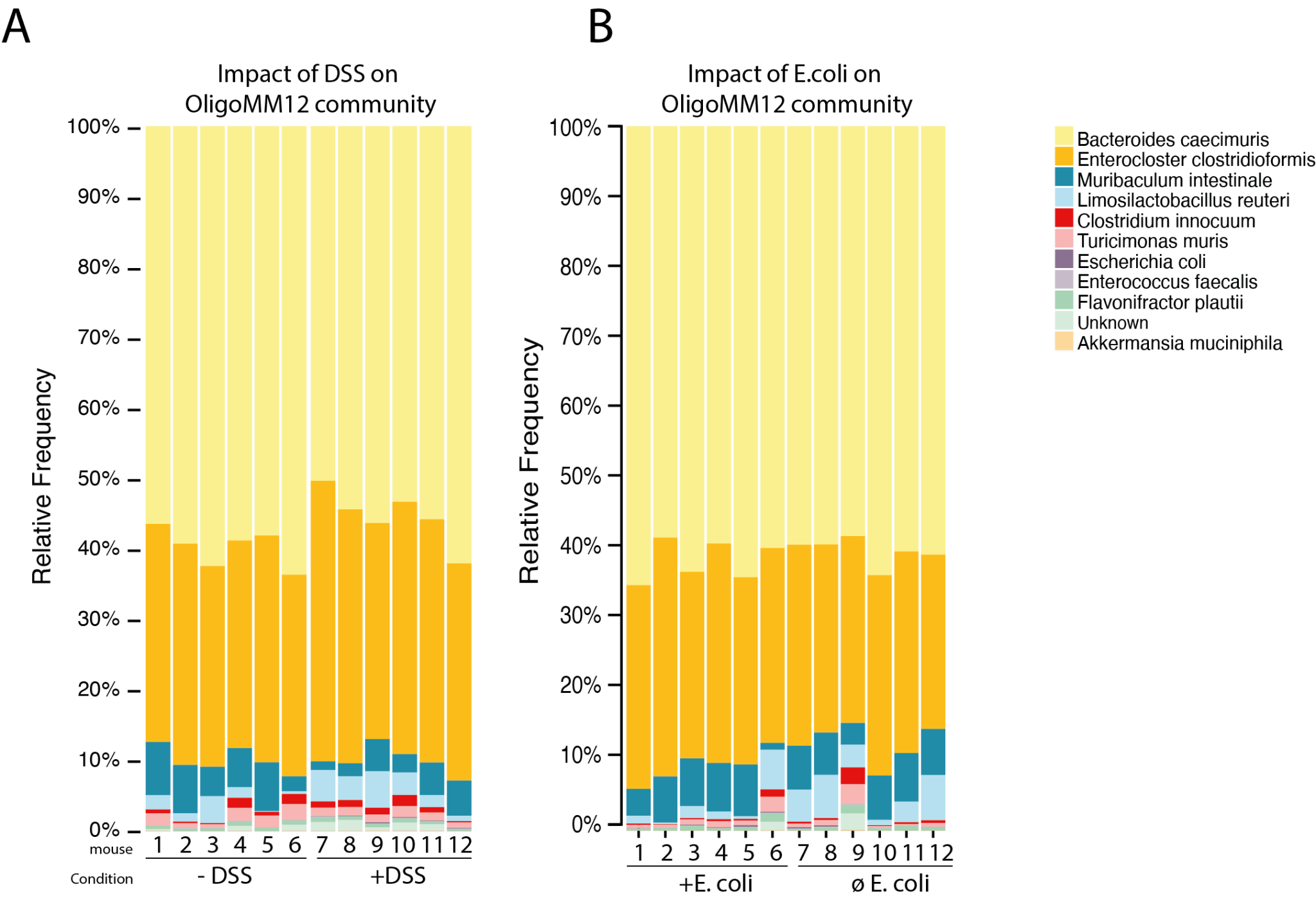
**Supplementary Figure 8. 16S rRNA gene analysis of the OligoMM12 community composition in feces under different conditions.**

**(A)** Comparison of community composition between H₂O- and DSS-treated mice.

**(B)** Comparison of community composition between mice colonized with *E. coli* and those without *E. coli* gavage.

In both cases, the results show that DSS-induced colitis and *E. coli* colonization do not significantly alter the overall composition or structure of the OligoMM12 community.

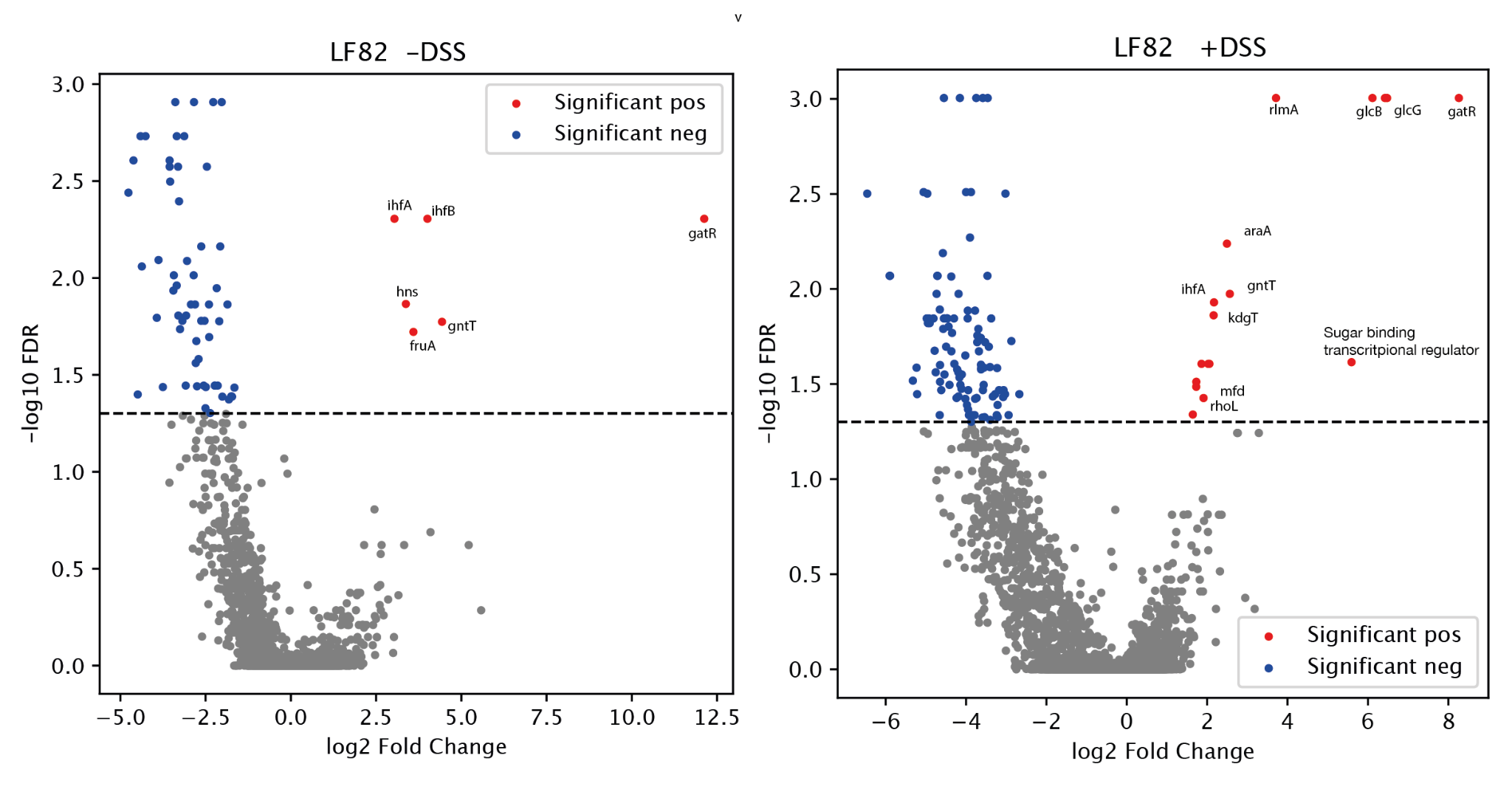

**Supplementary Figure 9. Volcano plot showing significantly essential and deleterious genes for LF82 in +DSS and non-DSS conditions**

-log10(FDR) obtained for each gene with Mageck analysis in function of the gene log2FC. Red genes are significantly deleterious, blue genes are significantly essential.

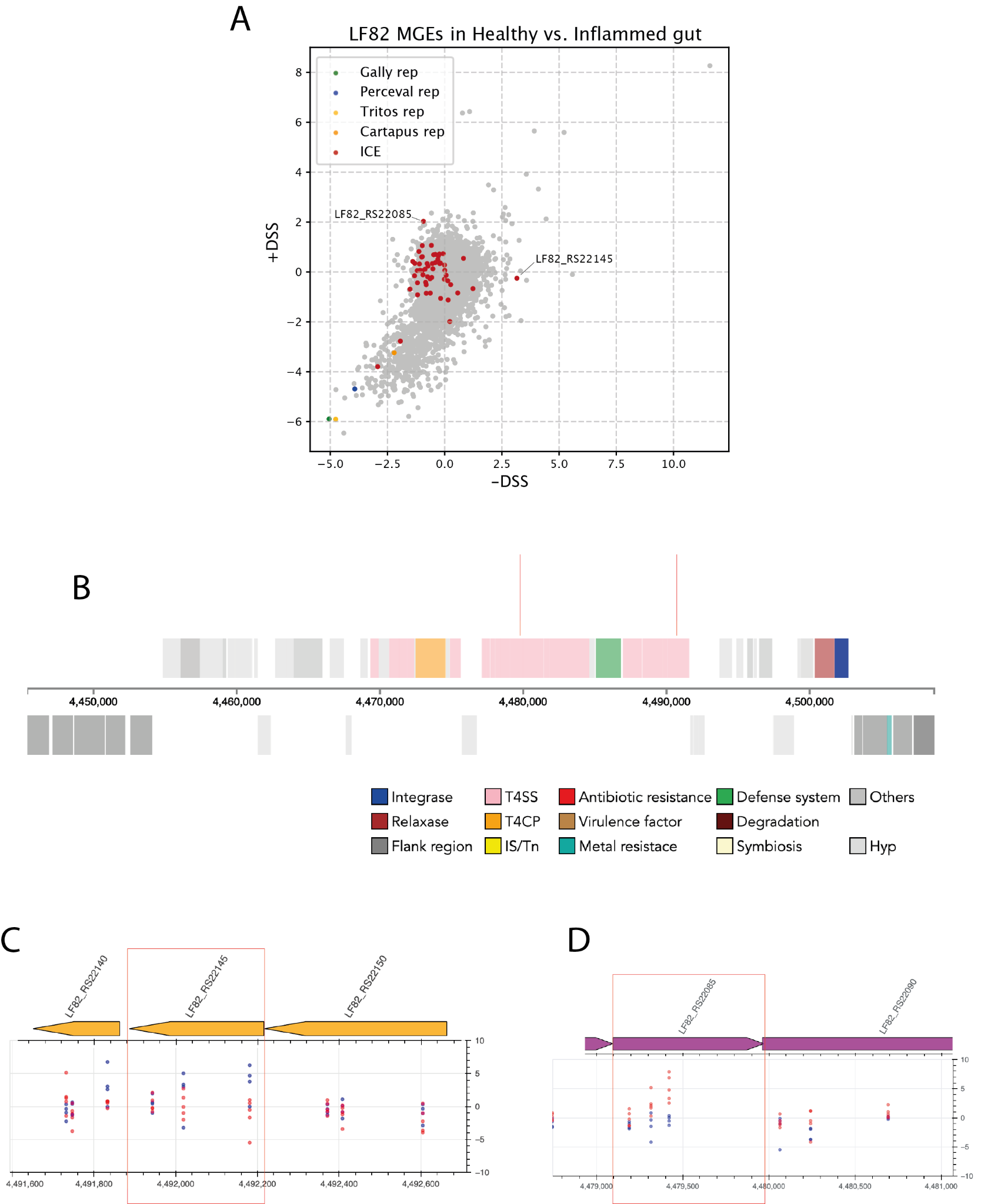

**Supplementary Figure 10. Mobile Genetic Elements in LF82**

**(C)** Scatter plot showing gene log₂FC values for LF82 in HO- *versus* DSS-treated mice. The four prophage repressors are highlighted in green, blue, yellow, and orange; integrative conjugative element (ICE) genes are shown in red. **(D)** LF82 ICE: Genome browser view zoomed in on genes belonging to the type IV secretion system (T4SS) and showing a strong differential signal between DSS- and H₂O-treated mice. Each dot represents a guide in either a H₂O-treated mouse (blue) or a DSS-treated mouse (red), with corresponding log₂FC values.
